# Acetylation-mediated phase control of the nucleolus regulates cellular acetyl-CoA responses

**DOI:** 10.1101/2020.01.24.918706

**Authors:** Ryan Houston, Shiori Sekine, Michael J. Calderon, Fayaz Seifuddin, Guanghui Wang, Hiroyuki Kawagishi, Daniela A. Malide, Yuesheng Li, Marjan Gucek, Mehdi Pirooznia, Alissa J. Nelson, Matthew P. Stokes, Jacob Stewart-Ornstein, Stacy G. Wendell, Simon C. Watkins, Toren Finkel, Yusuke Sekine

**Affiliations:** Aging Institute, Department of Medicine, University of Pittsburgh, Pittsburgh, PA, 15219 USA; Division of Cardiology, Department of Medicine, University of Pittsburgh, Pittsburgh, PA, 15213 USA; Department of Cell Biology, Center for Biologic Imaging, Univeristy of Pittsburgh, Pittsburgh, PA 15261, USA; National Heart, Lung, and Blood Institute, NIH, Bethesda, MD 20892, USA; Cell Signaling Technology, INC. Danvers, MA 01923, USA; Department of Computational and Systems Biology, University of Pittsburgh and Hillman Cancer Center, Pittsburgh, PA 15260, USA; Department of Pharmacology and Chemical Biology, University of Pittsburgh, Pittsburgh PA, 15261 USA; Division of Endocrinology and Metabolism, Department of Medicine, University of Pittsburgh, Pittsburgh PA, 15213 USA

**Keywords:** acetyl-CoA, acetylation, nucleolus, stress response, p53, Class IIa HDAC family, ribosomal protein, membraneless organelle, liquid-liquid phase separation

## Abstract

The metabolite acetyl-CoA serves as an essential element for a wide range of cellular functions including ATP production, lipid synthesis and protein acetylation. Intracellular acetyl-CoA concentrations are associated with nutrient availability, but the mechanisms by which a cell responds to fluctuations in acetyl-CoA levels remain elusive. Here, we generate a cell system to selectively manipulate the nucleo-cytoplasmic levels of acetyl-CoA using CRISPR-mediated gene editing and acetate supplementation of the culture media. Using this system and quantitative omics analyses, we demonstrate that acetyl-CoA depletion alters the integrity of the nucleolus, impairing ribosomal RNA synthesis and evoking the ribosomal protein-dependent activation of p53. This nucleolar remodeling appears to be mediated through the class IIa HDAC deacetylases regulating the phase state of the nucleolus. Our findings highlight acetylation-mediated control of the nucleolus as an important hub linking acetyl-CoA fluctuations to cellular stress responses.

## Introduction

Intracellular metabolites dynamically fluctuate in living organisms, in association with the availability of their source nutrients, such as glucose, lipids and amino acids. A cell employs a variety of molecular machineries to monitor the concentration of these metabolites. Upon metabolic fluctuations, those machineries activate signaling pathways that modulate gene expression and protein function to maintain metabolic homeostasis. Molecular links between the metabolite sensing and cellular responses have been extensively studied and several dedicated molecular networks have been described (Campbell and Wellen, 2018; Efeyan et al., 2015; Wang and Lei, 2018). However, partly due to experimental difficulties to selectively manipulate target metabolites among highly interconnected metabolic networks, the mechanisms for cellular primary responses towards fluctuations for most metabolites are not understood.

Acetyl-coenzyme A (acetyl-CoA) is a central metabolite that integrates diverse nutritional inputs into the biosynthesis of essential biomaterials including ATP, fatty acids and steroids, and therefore can be viewed as a critical indicator of the cellular metabolic state (Pietrocola et al., 2015; Shi and Tu, 2015). In addition, acetyl-CoA provides an acetyl donor for acetyltransferases catalyzing protein acetylation that modulates biophysical properties of modified proteins, thereby altering their protein stability, enzymatic activity, or interactions with other proteins. This modification is reversed by deacetylases. Hence, in accordance with the nutrient availability and the cellular metabolic state, protein acetylation can reversibly regulate a variety of biological processes including gene expression, signal transduction pathways and metabolic flux (Choudhary et al., 2014; Menzies et al., 2016).

In mammalian cells, acetyl-CoA is compartmentalized into a mitochondrial pool, and a nuclear/cytoplasmic pool (Pietrocola et al., 2015; Sivanand et al., 2018). In the mitochondrial matrix, acetyl-CoA is generated by the metabolism of nutrients including glucose, fatty acids and amino acids. Mitochondrial acetyl-CoA can enter the tricarboxylic acid (TCA) cycle thereby generating ATP and reducing equivalents (e.g. NADH) or it can be utilized for acetylation of mitochondrial proteins. Since acetyl-CoA is membrane impermeant, there is no direct exchange between mitochondrial acetyl-CoA and the acetyl-CoA in the cytosol. Rather, the TCA intermediate citrate can be exported from the mitochondria to the cytosol where it can become the predominant source for cytoplasmic and nuclear acetyl-CoA. The mitochondrial exported citrate is converted to acetyl-CoA and oxaloacetate by the enzyme ATP citrate lyase (ACLY) (Wellen et al., 2009). This nucleo-cytoplasmic acetyl-CoA pool serves as a building block for lipid synthesis including fatty acids, cholesterol and steroids, as well as serving as an acetyl donor for acetylation of cytosolic and nuclear proteins including histones. Another source of nucleo-cytoplasmic acetyl-CoA comes from the metabolite acetate, which derives from various extra- and intracellular sources such as food digestion, gut microbial metabolism, alcohol oxidation and deacetylation of acetylated proteins (Schug et al., 2016). Acetate can be utilized for acetyl-CoA synthesis through a reaction catalyzed by the enzyme acyl-CoA synthetase short chain family member 2 (ACSS2). In contrast to whole body *ACLY* deficient mice that are embryonic lethal, *ACSS2* deficient mice have no apparent phenotype under normal breeding conditions (Beigneux et al., 2004; Comerford et al., 2014). This would support a predominant role of the citrate-ACLY pathway for the nucleo-cytoplasmic acetyl-CoA production during development, and potentially under other circumstances as well. However, it should be noted that in some mouse tumor models or hypoxic tumor cells, or in other conditions where mitochondrial metabolism is dampened, the acetate-ACSS2 pathway can play a critical role in cell proliferation, lipid synthesis and histone acetylation (Comerford et al., 2014; Gao et al., 2016; Mashimo et al., 2014; Schug et al., 2015; Yoshii et al., 2009). Moreover, *ACLY* deficient mouse embryonic fibroblasts (MEFs) exhibit upregulation of ACSS2 and exogenously added acetate can be utilized for acetyl-CoA production in these cells (Zhao et al., 2016). These observations indicate a coordinated relationship between these two pathways in order to ensure the requisite supply of acetyl-CoA in the nucleo-cytoplasmic compartment.

Intimate connections between nucleo-cytoplasmic acetyl-CoA levels, the status of protein acetylation and cellular responses have been illustrated by multiple recent studies. In yeast cells during continuous glucose-limited growth, oscillations of acetyl-CoA are observed in accordance with distinct metabolic phases and an increase in acetyl-CoA is highly correlated with acetylation of several proteins including transcriptional coactivators and histones (Cai et al., 2011). Also, in various mammalian cell models, transcription is regulated either by acetyl-CoA abundance, or acetyl-CoA producing enzymes, i.e. ACLY or ACSS2, or by their nutrient sources (Gao et al., 2016; Huang et al., 2018; Lee et al., 2018; Lee et al., 2014; Li et al., 2017; Mews et al., 2017). Moreover, nutrient deprivation or starvation causes a rapid decline in acetyl-CoA abundance in cells in culture and in some mouse tissues, which is accompanied by deacetylation of proteins (Marino et al., 2014). In mammalian cells, as well as in yeast, pharmacological or genetic manipulations to deplete cytosolic acetyl-CoA have been reported to induce autophagy (Eisenberg et al., 2014; Marino et al., 2014). Moreover, the acetylation status of autophagy proteins, regulated by the p300 acetyltransferase and multiple deacetylases, plays a crucial role for autophagy induction (Lee et al., 2008; Lee and Finkel, 2009; Marino et al., 2014). As such, acetyl-CoA fluctuations appear to influence various biological responses through alterations in protein acetylation.

Here, we have developed a cell line which lacks ACLY and whose nucleo-cytoplasmic acetyl-CoA production is thereby solely dependent on exogenously supplemented acetate. Removal of acetate from the culture media enables us to rapidly deplete intracellular acetyl-CoA. Using this cell line, we provide quantitative data sets for alterations in mRNA levels, protein abundance and acetylation status in cells experiencing acute acetyl-CoA depletion. These analyses uncovered that in response to acetyl-CoA depletion, the integrity of the nucleolus, an organelle critical for ribosomal biosynthesis, was morphologically, biophysically and functionally remodeled. This nucleolus remodeling leads to the upregulation of the stress responsive transcription factor p53 in a ribosomal protein-dependent manner. Furthermore, these alterations were found to be suppressed by class IIa HDAC inhibitors. We identified multiple nucleolar proteins whose acetylation levels were potentially regulated by the class IIa HDACs, suggesting that class IIa HDAC-dependent deacetylation of nucleolar proteins may play an important role in regulating the integrity of the nucleolus and the induction of a nucleolus-dependent stress response when acetyl-CoA levels decline.

## Results

### Acetate-dependent control of acetyl-CoA production in ACLY KO cells

In order to understand cellular responses to fluctuations in nucleo-cytoplasmic acetyl-CoA abundances, we sought to devise a cell system where we could selectively and robustly control acetyl-CoA levels. To achieve this, we focused on establishing a cell line whose acetyl-CoA in the nucleo-cytoplasmic compartment was solely synthesized through the acetate-ACSS2 axis, assuming that such a cell line would enable us to manipulate acetyl-CoA levels by simply modulating the amount of acetate exogenously supplemented in the culture media (**Figure 1A**). Hence, using the CRISPR-Cas9 system, we targeted the *ACLY* gene in HT1080 human fibrosarcoma cells to make *ACLY* deficient cell lines. A plasmid coding an *ACLY*-targeting single guide RNA (sgRNA) together with Cas9 was transfected into HT1080 cells and the transfected cell clones were recovered either in the standard culture media (0 mM added acetate) or in the same media but with excess amounts (2 or 20 mM) of sodium acetate (**Figure 1-figure supplement 1A**). Among the cell clones screened, putative *ACLY* genome-edited clones were found to only exist in the acetate supplemented media (**Figure 1-figure supplement 1B**), suggesting that acetate supplementation increased the viability or growth of *ACLY* deficient cells. This observation was consistent with the previously reported phenotype of *ACLY* deficient MEFs (Zhao et al., 2016). We selected two independent *ACLY* deficient (KO) and ACLY expressing (WT) clones, respectively, that henceforth were continuously cultured in the presence of 20 mM acetate for further analyses (**Figure 1B and Figure 1-figure supplement 1C**). Hereafter, we refer to these cells as ASA-KO and ASA-WT (<u>a</u>cetate-<u>s</u>upplemented *<u>A</u>CLY* <u>KO</u> and <u>WT</u>), respectively. In this culture condition, ASA-KO cells did not exhibit any defect in viability when compared with ASA-WT cells (**Figure 1C**, (+) Acetate, 10% FBS). However, removal of acetate from the culture media for 4 days significantly impaired the viability of ASA-KO but not ASA-WT cells (**Figure 1C**). Consistent with the observation that fetal bovine serum (FBS) contains submillimolar concentration of acetate (Kamphorst et al., 2014; Zhao et al., 2016), either a decrease in FBS concentration from 10% to 1% or the use of 10% dialyzed FBS (dFBS) enhanced the vulnerability of ASA-KO cells, while these manipulations had no significant effect on viability in the acetate-supplemented conditions (**Figure 1C**). Moreover, exogenous expression of ACLY in ASA-KO cells rescued the sensitivity to acetate removal, indicating that the acetate dependence of these cells was caused by *ACLY* deficiency (**Figure 1D**).

**Figure 1.**
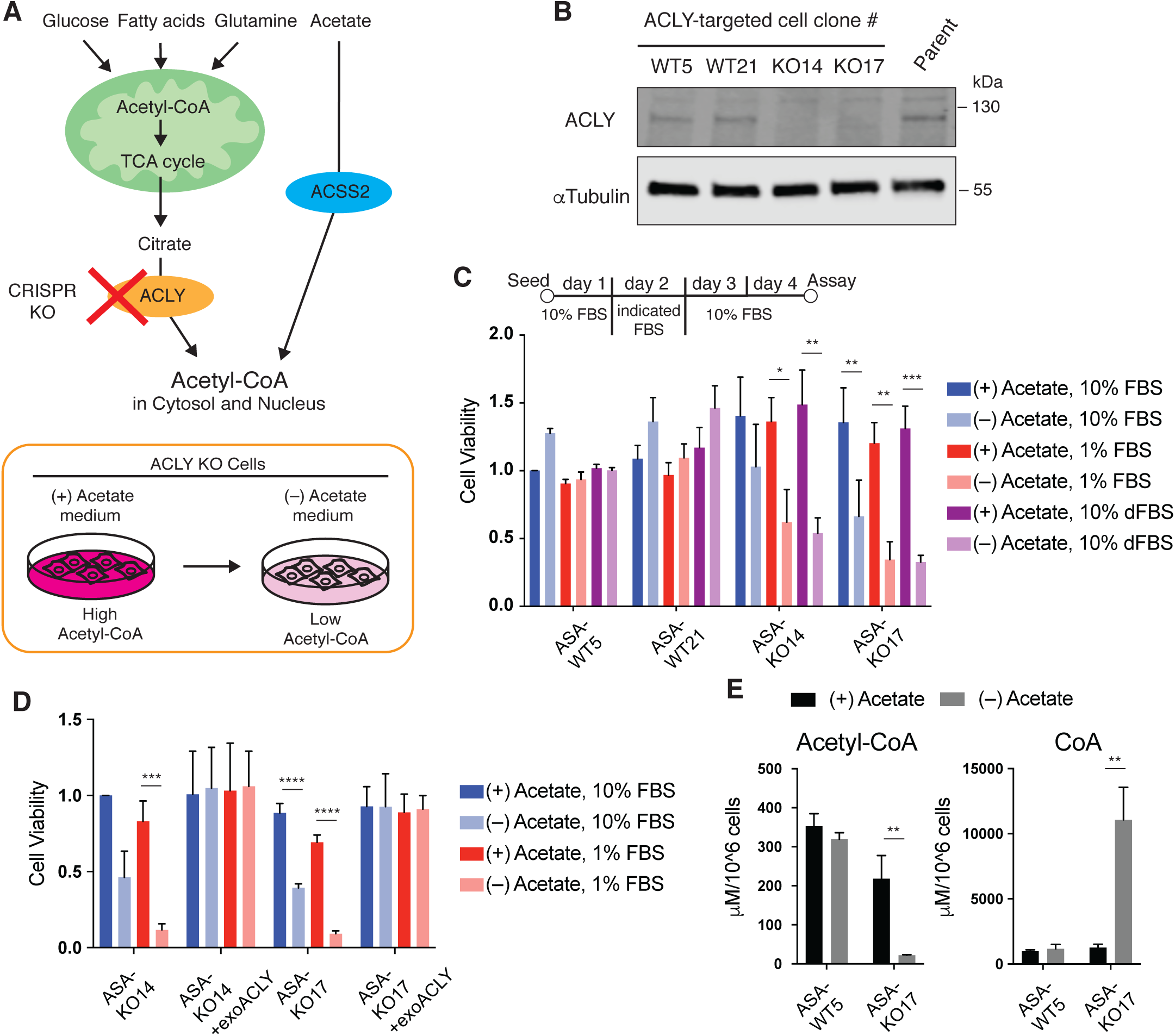
Acetate dependency for acetyl-CoA production in ASA-KO cells. (A) Schematic representation of the pathways for nucleo-cytoplasmic acetyl-CoA production in mammalian cells (Top) and of the acetate-dependent control of acetyl-CoA in ACLY KO cells (bottom). (B) Immunoblotting for ACLY in representative ACLY-targeted cell clones maintained in 20 mM acetate-supplemented media and in parental HT1080 cells. *a*Tubulin is shown as a loading control. (C) Cell viability assay for ASA-WT and KO cells cultured in the presence or absence of 20 mM acetate in the culture media with indicated FBS conditions over 4 days. On day 2, the original medium containing 10% FBS was replaced for 24 hours with media containing either 10% FBS, 1% FBS, or 10% dialyzed FBS (dFBS). Data are shown as mean ± S.D. (n=3 independent experiments). *P<0.05, **P<0.01, ***P<0.001 (One-way ANOVA followed by Tukey multiple comparisons test). (D) Cell viability assay for ASA-KO cells with or without exogenous expression of ACLY (exoACLY) using the same experimental procedures as in “C”. Data are shown as mean ± S.D. (n=3 independent experiments). ***P<0.001, ****P<0.0001 (One-way ANOVA followed by Tukey multiple comparisons test). (E) Quantification of acetyl-CoA and Coenzyme A (CoA) in cell lysate from ASA-WT or KO cells cultured for 4 hours in media containing 1% FBS with or without 20 mM acetate. Data are shown as mean ± S.D. (n=3 biological replicates). **P<0.01 (Unpaired t test).

Using this cell system, we next examined whether acetate removal modulated intracellular acetyl-CoA levels in ASA-KO cells. We used media containing 1% FBS during acetate removal to maximize the effect of acetate withdrawal, although similar, albeit slightly milder effects were seen following acetate withdrawal in the setting of 10% FBS. Quantifications of acetyl-CoA in the whole cell lysates demonstrated that withdrawing acetate for 4 hours drastically decreased the amount of acetyl-CoA in ASA-KO cells, while this intervention had only a marginal effect in ASA-WT cells (**Figure 1E**). In contrast, the amount of Coenzyme A (CoA) was reciprocally increased in ASA-KO cells in response to acetate removal (**Figure 1E**). Taken together, in ASA-KO cells, even in the presence of other nutrient sources such as glucose and lipids, the nucleo-cytoplasic acetyl-CoA levels are seemingly tunable solely by adding or removing acetate in the culture media.

### Acetyl-CoA depletion modulates protein acetylation

Using the ASA-KO cell system, we profiled global cellular responses following acute acetyl-CoA depletion. As cytoplasmic acetyl-CoA is mainly utilized for lipid synthesis and protein acetylation, we examined whether a rapid decline in acetyl-CoA levels affected these downstream events. Quantifications of total cholesterol and a series of fatty acids revealed that although some fatty acids including linoleic acid and Dihomo-*γ*-linolenic acid (DGLA) exhibited significant decreases, overall alterations in total amounts of cholesterol and fatty acids after 4 hours of acetate removal were modest when compared to the dramatic decline in acetyl-CoA levels at the same time point (**Figure 2A** and **Figure 1E**). Thus, at least at early time points, dramatic reductions in acetyl-CoA are not fully reflected by marked alterations in total cholesterol or lipid amounts.

**Figure 2.**
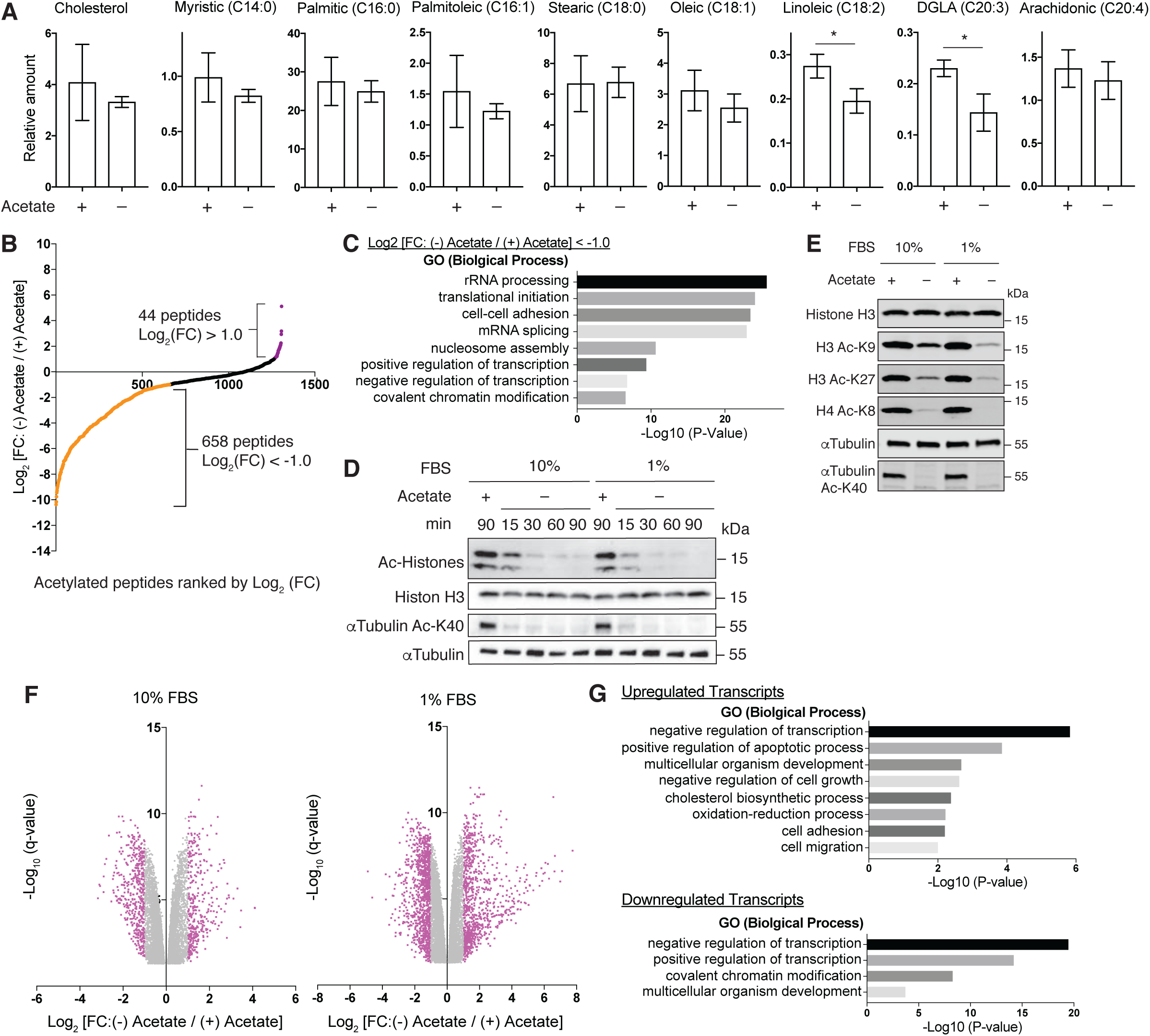
Alterations in lipid synthesis, acetylation and gene expression following acetyl-CoA depletion. (A) Quantification of total cholesterol and fatty acids in ASA-KO cells cultured for 4 hours in 1% FBS containing media with or without 20 mM acetate. Data are shown as mean of 3 biological replicates. *P<0.05 (Unpaired t test). (B) Plot of acetylated peptides ranked by log_2_ fold changes 90 minutes after acetate removal in ASA-KO cells. Values are from Table S1, Column B (mean of two independent technical replicates). (C) Gene ontology (GO) analysis for putative deacetylated peptides identified in the acetylome analysis. Enriched representative Biological Process are shown. (D) Immunoblotting for the indicated acetylated proteins in ASA-KO cells cultured in 10% or 1% FBS containing media with or without 20 mM acetate for the indicated times. Acetylated (Ac) histones were detected using an anti-acetyl-lysine antibody. (E) Immunoblotting for the designated acetylated and total protein level in ASA-KO cells cultured in 10% or 1% FBS containing media with or without acetate for 4 hours. (F) Volcano plots for mRNA expression changes 4 hours after acetate removal in ASA-KO cells cultured in 10% or 1% FBS containing media. Data shown represents the mean of 3 biological replicates. Values are from Table S2 and genes with P<0.05 are shown. (G) Gene ontology (GO) analysis for downregulated transcripts (log_2_ FC [(-) Acetate / (+) Acetate] < −1.0. q<0.05) and upregulated transcripts (log_2_ FC [(-) Acetate / (+) Acetate] >1.0. q<0.05) in the RNA sequencing with the 1% FBS condition. Enriched representative Biological Process are shown.

In order to examine global changes in the status of protein acetylation upon acetyl-CoA depletion, we conducted an acetylome analysis using an acetyl-lysine motif antibody-based immunoaffinity purification. Recovered acetylated lysine-containing peptides were then detected through liquid chromatography mass-spectrometry. This approach identified 1307 acetylated lysine-containing peptides in ASA-KO cells cultured in the presence of acetate (**Table S1**). Among these, 658 peptides exhibited more than a 50% decrease after 90 minutes of acetate withdrawal {log_2_ [fold change (FC): (-) Acetate / (+) Acetate] < −1.0}, suggesting that acetyl-CoA depletion caused rapid deacetylation of nearly half of all acetylated proteins (**Figure 2B and Table S1**, Column B). These peptides included a wide variety of proteins including molecules involved in transcription, translation, ribosomal RNA (rRNA) processing, messenger RNA (mRNA) splicing, nucleosome assembly and cell-cell adherence junctions (**Figure 2C**). By immunoblotting using pan- and site specific-anti-acetyl-lysine antibodies, we detected immediate and robust decreases in acetylation of both histone and non-histone proteins upon acetate withdrawal (**Figure 2D-2E**). Collectively, these findings suggest that unlike the cholesterol or lipid pool, alterations in intracellular acetyl-CoA levels are rapidly reflected in the status of protein acetylation.

### Acetyl-CoA depletion modulates gene expressions

Since we observed marked alterations in protein acetylation including on histones, we performed transcriptome analyses by RNA sequencing of ASA-KO cells cultured in the presence or absence of acetate to examine the transcriptional alterations following acetyl-CoA depletion. Consistent with a notion that histone deacetylation suppresses transcription, we identified a large number of transcripts downregulated 4 hours after acetate withdrawal {526 and 1096 transcripts in 10% FBS and 1% FBS conditions, respectively, log_2_ [FC: (-) Acetate / (+) Acetate] < −1.0, q < 0.05} (**Figure 2F and Table S2**, sheet 1 and sheet 2 for 10% FBS and 1% FBS conditions, respectively). Intriguingly, we also found many transcripts that were significantly upregulated in these conditions {365 and 899 transcripts in 10% FBS and 1% FBS conditions, respectively, log_2_ [FC: (-) Acetate / (+) Acetate] > 1.0, q < 0.05}. These differentially expressed transcripts in the 10% and 1% FBS contidions were highly overlapped (70.5% or 33.8% of downregulated transcritps in 10% or 1% FBS conditions, respectively, and 65.2% or 26.4% of upregulated transcritps in 10% or 1% FBS conditions, respectively, **Figure 2-figure supplement 1A**), suggesting that a smillar but stronger transcriptional response occurs in the 1% FBS condition. Gene ontology analyses for differentially expressed genes in the 1% FBS condition uncovered that several clusters of genes were similarly regulated (**Figure 2G**). For example, genes related to transcriptional regulation were highly enriched in both up- and down-regulated transcripts. Of particular interest was that genes involved in cellular stress responses such as apopototic process, cell cycle arrest and oxidation-reduction process were selectively upregulated in acetate-withdrawn cells, suggesting that stress signaling pathways are activated in response to acetyl-CoA depeltion.

### Acetyl-CoA depletion alters the integrity of the nucleolus

To further understand the complete cellular response after acetyl-CoA depletion, we also performed quantitative proteomics analyses using a tandem mass tag (TMT) system with the same experimental conditions used for our previous experiments (i.e. analysis 4 hours after acetate removal in 1% FBS containing media). Although overall alterations in protein levels appeared to be less drastic, we observed that the expression of 51 proteins were significantly increased more than 1.2-fold and 135 proteins were decreased less than 0.8-fold in response to acetate removal (**Table S3**). Intriguingly, we found that many proteins that are known to be localized to the nucleolus were significantly altered (**Figure 3A**, shown in red). Changes in these nucleolar proteins were not evident on the RNA seq analysis (**Figure 3A**, y-axis), implying that these proteins were likely regulated at a post-transcriptional level. By immunoblotting, we confirmed that ribosome biogenesis factors RRP1 and BOP1 were increased while PICT1 (also known as NOP53 or GLTSCR2) was decreased after acetate removal in ASA-KO cells but not in ASA-WT cells (**Figure 3-figure supplement 1A and 1B**). These observations indicate that alterations of the nucleolus might occur upon acetyl-CoA depletion. Hence, we next addressed whether acetyl-CoA depletion affected nucleolar structures and functions. The nucleolus is the organelle primarily responsible for ribosomal biosynthesis, where rRNA synthesis, rRNA processing and assembly of rRNA and ribosomal proteins takes place (Mangan et al., 2017; Nemeth and Grummt, 2018). Using a 5-Fluorouridine (FUrd) incorporation assay, we monitored newly synthesized rRNA in the nucleolus and found that acetate removal rapidly reduced rRNA synthesis in ASA-KO but not ASA-WT cells (**Figure 3B-3C**). We also utilized a rRNA specific dye to stain cells in conjunction with immunostaining for nucleolar proteins. In the presence of acetate, the nucleolar transcription factor UBF localized within the nucleolar region as evident by its overlap with rRNA (**Figure 3D**). Consistent with observed the impairment of rRNA synthesis (**Figure 3B-3C**), fluorescent intensities for nuclear rRNA were significantly decreased in ASA-KO cells upon acetate removal (**Figure 3D-3E**). Interestingly, residual rRNA signals detected in the acetate-deprived ASA-KO cells seemingly segregated from UBF (**Figure 3D**, magnified nuclear images with the line profiles). The nucleolar structure can be divided into three distinct functional subcompartments including the UBF containing compartment for rRNA synthesis, as well as compartments for rRNA splicing or for rRNA-ribosomal protein assembry (**Figure 3-figure supplement 1C**) (Thiry and Lafontaine, 2005). We therefore performed a co-staining of rRNA with marker proteins for the latter two compartments; fibrilalin (FBL) or nucleolin (NCL), respectively. Interestingly, while both proteins colocalized with rRNA in the acetate-supplemented cells, neither FBL nor NCL fully merged with the segregated rRNA signals in the acetate-deprived ASA-KO cells (**Figure 3-figure supplement 1D-1E**). These observations imply that acetyl-CoA depletion redistributes the rRNA containing compartment within the nucleolus. Furthermore, we found that in ASA-KO cells, the nucleolar localization of a nucleolar scaffolding protein, nucleophosmin (NPM1), became dispersed following acetate removal (**Figure 3F**). We also noted that ASA-WT cells did not exhibit any of these morphological alterations in the nucleolus in response to acetate removal (**Figure 3-figure supplement 1F-1G**). Altogether, these observations suggest that acetyl-CoA depletion markedly influences nucleolar components including changes in levels and in localization, thereby structually and functionally remodeling the nucleolus.

**Figure 3.**
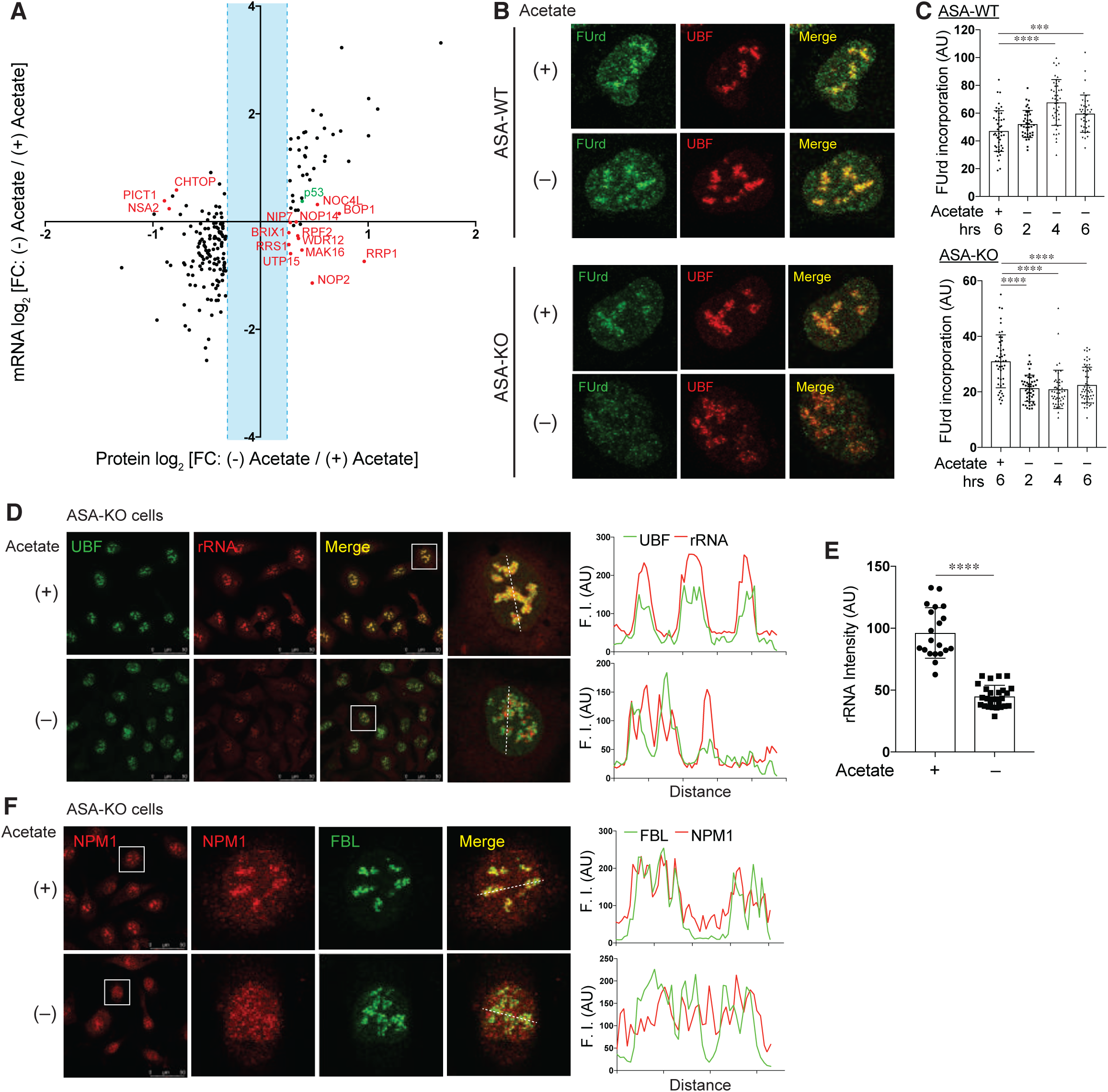
Functional and structural remodeling of the nucleolus by acetyl-CoA depletion. (A) Scatter plot for log_2_ fold changes (FC) in proteins (X-axis) and in mRNA (Y-axis) 4 hours after acetate removal in ASA-KO cells. Proteins with log_2_ fold changes more than 0.264 or less than −0.322 are shown. Proteins that are known to localize in the nucleolus are highlighted in red and p53 shown in green. (B) FUrd incorporation assay in ASA-WT or ASA-KO cells cultured in 1% FBS containing media with or without 20 mM acetate over 6 hours. Representative nuclear images immunostained for FUrd and UBF are shown. (C) Quantification of the mean fluorescent intensity of the FUrd signal per nucleus. The microscopic images of the FUrd incorporation assay performed as in (B) were utilized for the quantification. Data are shown as mean ± S.D. (n=39-55 cells per condition). ***P<0.001, ****P<0.0001 (One-way ANOVA followed by Tukey multiple comparisons test). AU, arbitrary unit. (D) Immunostaining for UBF along with rRNA dye staining in ASA-KO cells cultured in 1% FBS containing media with or without acetate for 4 hours. Magnified nuclear images (surrounded by a white square) are shown. Line profiles for indicated fluorescent intensities (F. I) determined along the white dashed lines are shown to the right. (E) Quantification of the mean fluorescent intensity of the rRNA signal per nucleus in (D). Data are shown as mean ± S.D. [n=20 or 27 cells for (+) Acetate or (-) Acetate, respectively]. ****P<0.0001 (Unpaired t test). (F) Immunostaining for NPM1 and FBL in ASA-KO cells cultured in 1% FBS containing media with or without acetate for 4 hours. Magnified nuclear images (surrounded by a white square) are shown. Line profiles for indicated fluorescent intensities (F. I) determined along the white dashed lines are shown to the right.

### Acetyl-CoA depletion induces ribosomal protein-dependent p53 activation

In addition to the nucleolar alterations, we observed an increase in the level of p53 protein in the quantitative proteomics of acetyl-CoA-depleted cells (**Figure 3A**, shown in green). We also found upregulation of the p53 target genes, including *MDM2*, *CDKN1A, BBC3* and *PLK3*, in our RNA sequencing analysis (**Figure 4-figure supplement 1A**). We therefore next demonstrated by immunoblotting that removing acetate from the media increased p53 expression only in ASA-KO cells (**Figure 4A**). A number of studies have demonstrated that so called “nucleolar stress”, which is caused by aberrations in ribosome biogenesis, induces p53 stabilization in a ribosomal protein-dependent manner (Boulon et al., 2010; Deisenroth et al., 2016). Mechanistically, the most well studied ribosomal proteins in this context are RPL11 and RPL5. These components are incorporated into a large ribosomal subunit by forming a complex with 5S rRNA in the unstressed nucleolus. Once the nucleolus is stressed by, for instance, the RNA pol I inhibitor actinomycin D (ActD), the 5S-rRNA-RPL11-RPL5 complex is known to translocate to the nucleoplasm, where it interacts and inhibits MDM2, an E3 ubiquitin ligase for p53, thereby stabilizing p53 (**Figure 4B**) (Donati et al., 2013; Sloan et al., 2013). Moreover, it has been reported that PICT1 is degraded by nucleolar stressors and that depletion of PICT1 is sufficient for p53 induction in a ribosomal protein-dependent manner (Maehama et al., 2014; Sasaki et al., 2011). Since we observed that acetyl-CoA depletion induced nucleolar alterations and reciprocal alterations in p53 and PICT1 levels, we next assessed whether these responses were mediated through a nucleolar stress condition. Upon acetate removal, interactions between endogenous MDM2 and FLAG-tagged RPL11 were found to increase in a time-dependent manner in ASA-KO cells (**Figure 4C**). Moreover, small interfering (si) RNA-mediated knockdown of RPL11 or RPL5 diminished acetate-induced p53 upregulation, as well as PICT1 degradation, suggesting that RPL11 and RPL5 are required for these processes (**Figure 4D**). These observations were consistent with other nucleolar stress-dependent p53 activation (Donati et al., 2013; Sloan et al., 2013). DNA damage is a potent inducer of p53, but neither acetate removal nor ActD treatment induce phosphorylation of Serine 139 in Histone H2A.X (*γ*H2A.X), a marker for the DNA double strand breaks, suggesting DNA damage-independent mechanisms for the p53 upregulation observed in acetyl-CoA-depleted cells (**Figure 4-figure supplement 1B-1C**). Collectively, these observations suggest that acetyl-CoA depletion-induced p53 activation is mediated through ribosomal proteins associated with the nucleolar stress response.

**Figure 4.**
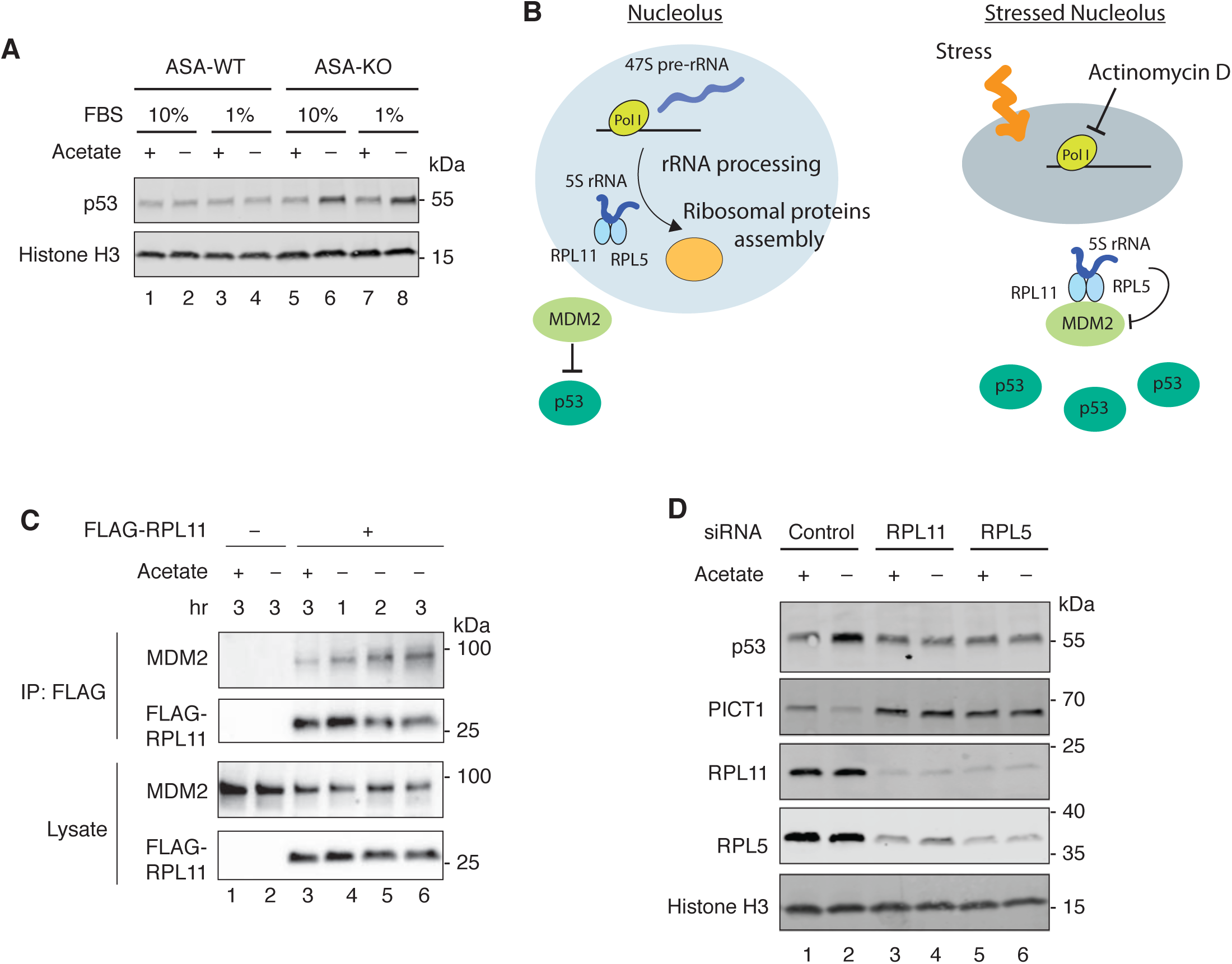
Ribosomal protein-dependent p53 activation by acetyl-CoA depletion. (A) Immunoblotting for p53 in ASA-WT and ASA-KO cells cultured in 10% or 1% FBS containing media with or without acetate for 4 hours. Histone H3 is shown as a loading control. (B) Schematic representation of the nucleolus as an organelle for ribosomal biogenesis (left) and the stressed nucleolus that induces p53 stabilization through the inhibitory interaction between the 5S rRNA-RPL11-RPL5 and MDM2. (C) Immunoprecipitation with FLAG-RPL11 and Immunoblotting for MDM2 and FLAG-RPL11 in ASA-KO cells cultured in 1% FBS containing media with or without acetate for the indicated hours. (D) Immunoblotting for p53 levels in ASA-KO cells pretreated with indicated siRNAs and cultured in 1% FBS containing media with or without acetate for 4 hours. Histone H3 is shown as a loading control.

### Class IIa HDAC family mediates acetyl-CoA-dependent nucleolar alterations

We then addressed what mediates these nucleolus-dependent responses induced by acetyl-CoA depletion. As described above, we found drastic alterations in protein acetylation in acetyl-CoA-depleted ASA-KO cells (**Figure 2B-2E and Table S1**). Therefore, we tested a possible role for lysine deacetylation in the acetyl-CoA depletion-dependent nucleolar alterations and subsequent p53 activation. The histone deacetylase (HDAC) family members are the predominant deacetylases in mammalian cells, which can be grouped into zinc ion-dependent (class I, II and IV) and nicotinamide adenine dinucleotide (NAD^+^)-dependent (class III, also known as Sirtuin family) families (**Figure 5A**) (Haberland et al., 2009). We utilized chemical inibitors for HDACs as this way enabled us to transiently inhibit HDACs while depleting acetate. We demonstrated that general inhibitors for zinc-dependent HDACs, Tricostatin A and Vorinostat (also known as SAHA) completely inhibited the acetyl-CoA depletion-induced reciprocal regulation of p53 and PICT1 (**Figure 5B**), suggesting that zinc-dependent HDACs are likely required for this process. In contrast, EX-527, an inhibitor for the NAD^+^-dependent deacetylase Sirtuin 1, did not alter this response (**Figure 5-figure supplement 1A**). Using selective inhibitors for each HDAC class, we sought to further address which HDAC is responsible for the acetyl-CoA depletion-induced p53 and PICT1 regulation (**Figure 5A**). While the class I inhibitor Entinostat (also known as MS-275) and the class IIa inhibitor TMP195 exhibited differential inhibitory effect on deacetylation of Histone H3 K9 and *α*Tubulin K40, we found that TMP195 but not Entinostat blocked the reciprocal regulation of p53 and PICT1 (**Figure 5B**) (Lobera et al., 2013; Saito et al., 1999). TMP195 also suppressed the acetate-dependent induction of p21(CDKN1A), a downstream target of p53 (**Figure 5C**). Another class IIa inhibitor LMK235 also exhibited the same effect, while other class I inhibitors including Mocetinostat and RGFP966 (HDAC3 selective) or the class IIb HDAC6 inhibitor Tubastatin A did not (**Figure 5C** and **Figure 5-figure supplement 1B**) (Butler et al., 2010; Fournel et al., 2008; Malvaez et al., 2013). These observations let us hypothesize that the class IIa HDACs-mediated deacetylation may modulate the nucleolus. Thus, we investigated whether the inhibition of class IIa HDACs also maintained nucleolar integrity and rRNA synthesis using immunostaining with the rRNA dye and FUrd assay, respectively. We noted that the class IIa HDAC inhibitors TMP195 and LMK235 both restored the intensity of rRNA and its co-localization with NCL in acetyl-CoA-depleted ASA-KO cells (**Figure 5-figure supplement 5C-5D**). Furthermore, TMP195 and LMK235 recovered FUrd incorporation while the class I inhibitor Entinostat was ineffective (**Figure 5D-5E**). These findings suggest that the class IIa HDACs are selectively required for acetyl-CoA depletion-induced nucleolar remodeling and p53 activation.

**Figure 5.**
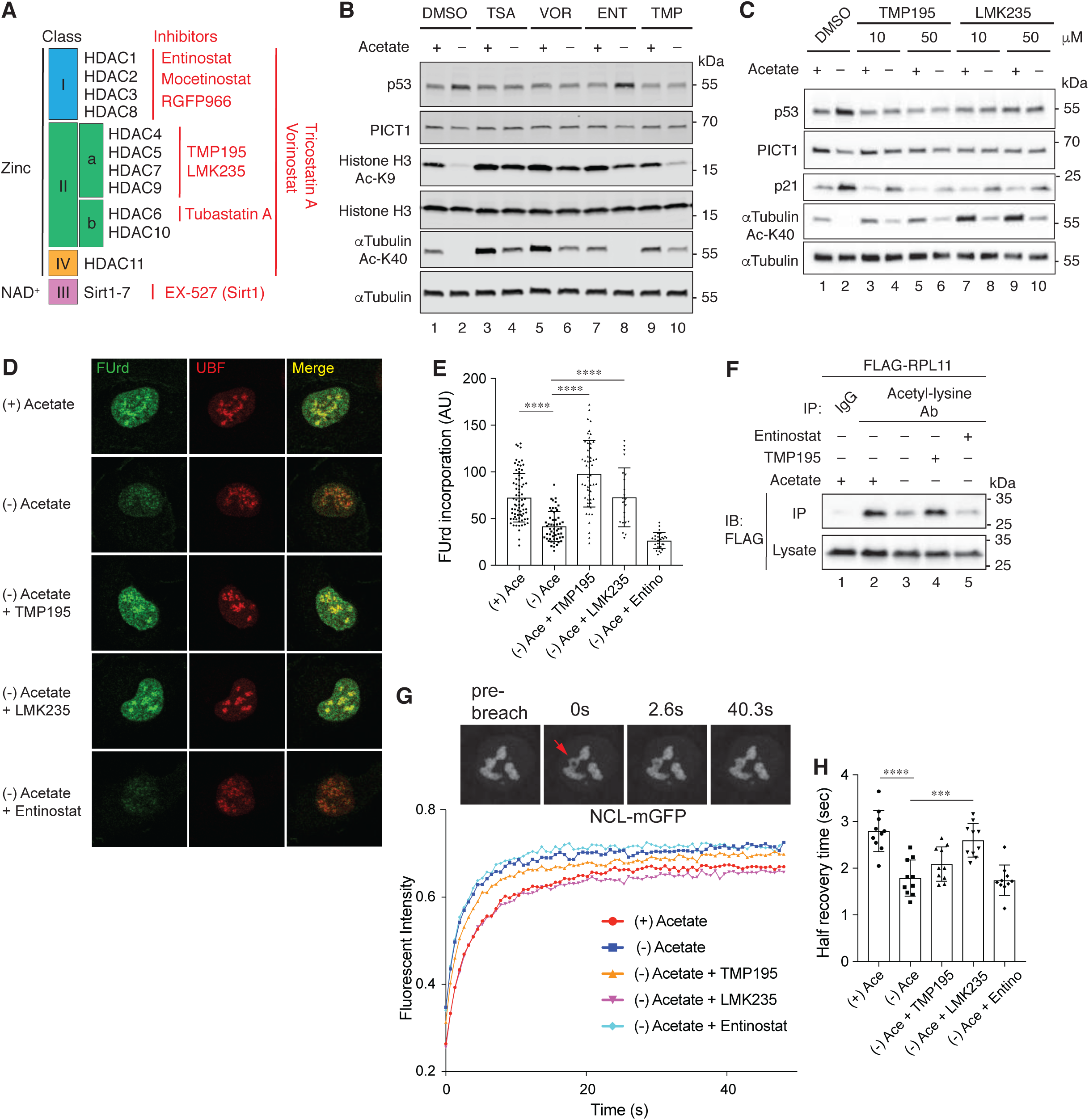
Class IIa HDAC activity is required for acetyl-CoA-induced nucleolar alterations. (A) Schematic representation of mammalian HDAC family members and their inhibitors (red). (B) Immunoblotting for the indicated proteins in ASA-KO cells cultured in 1% FBS containing media with or without acetate, and in the presence or absence of indicated HDAC inhibitors for 4 hours. Following concentrations of HDAC inhibitors were used: 500 nM Triconstatin A, 10 mM Vorinostat, 70 mM Entinostat, and 50 mM TMP195. (C) Immunoblotting for the indicated proteins in ASA-KO cells cultured in 1% FBS containing media with or without acetate, and in the presence or absence of indicated class IIa HDAC inhibitors for 4 hours. (D) FUrd incorporation assay in ASA-KO cells cultured in 1% FBS conditions with or without acetate and and in the presence or absence of indicated HDAC inhibitors for 4 hours. Representative nuclear images (surrounded by a white square) immunostainined for FUrd and UBF are shown. (E) Quantification of the mean fluorescent intensity of the FUrd signal per nucleus in (E). Data are shown as mean ± S.D. (n=23-74 cells per condition). ****P<0.0001 (One-way ANOVA followed by Tukey multiple comparisons test). AU, arbitrary unit. (F) Immunoprecipitation with an acetyl-lysine motif antibody and immunoblotting for FLAG-RPL11 in ASA-KO cells cultured in 1% FBS containing media with or without acetate, and in the presence or absence of 50 mM TMP195 or 50 mM M Entinostat for 90 minutes. (G) FRAP of NCL-mGFP in ASA-KO cells cultured in 1% FBS containing media with or without acetate, and in the presence or absence of 50 mM TMP195, 10 mM LMK235 or 50 mM Entinostat for 90 minutes. Data are shown as mean (n=10 independent cells from two independent experiments). Representative nucleolar images expressing NCL-mGFP before and after photobreaching are shown on the top. The red allow indicates the ROI for photobreaching. (H) Half recovery times (sec) obtained from the FRAP curves in “G”. Data are shown as mean ± S.D. (n=10 independent cells). ***P<0.001 ****P<0.0001 (One-way ANOVA followed by Tukey multiple comparisons test).

### Class IIa HDACs regulate deacetylation of nucleolar proteins and nucleolar dynamics

To investigate whether the effect of class IIa HDAC inhibitors on the nucleolus is mediated through inhibition of deacetylation, we conducted an acetylome analysis for acetyl-CoA-depleted cells with and without the class IIa inhibitor TMP195 and identified 365 acetylated peptides whose acetylation levels were maintained at a level two-fold or greater by TMP195 treatment in the setting of acetate removal {**Table S1**, Column C, Log_2_ [FC: (-) Acetate + TMP195 / (-) Acetate] >1.0}. Importantly, among these TMP195 sensitive proteins, we identified multiple nucleolus-resident proteins including ribosomal proteins and NCL (**Figure 5-figure supplement 2A**). Using the acetyl-lysine motif antibody immunoprecipitation assay, we demonstrated that acetylation of RPL11 decreased upon acetate removal, and that this was suppressed by TMP195 but not by the class I inhibitor Entinostat (**Figure 5F**). Deacetylation of NCL by acetate removal was also partially recovered by TMP195 (**Figure 5-figure supplement 2B**). These results suggest that the class IIa HDACs mediate deacetylation of at least certain nucleolar proteins including RPL11 and NCL.

The nucleolus is known to be formed through a process called liquid-liquid phase separation (LLPS), a central mechanism for facilitating the formation of biomolecular condensates that exhibit liquid droplet-like or other related material properties (Banani et al., 2017; Shin and Brangwynne, 2017; Snead and Gladfelter, 2019). Because LLPS is mediated through multivalent weak interactions among the component proteins and nucleotides, post translational modifications (PTMs) of proteins including phosphorylation, methylation and acetylation can shift the assembly and material properties of condenstates by modulating their interactions (Hofweber and Dormann, 2019; Snead and Gladfelter, 2019). Therefore, we sought to examine the effect of acetylation on the LLPS state of the nucleolus. As the molecular dynamics in a condensate can be seen as an indicator for its material property, we measured the mobility of NCL-mGFP by using fluorescence recovery after photobleaching (FRAP) (**Figure 5G**). We set a photobleaching area within the nucleoluar region marked by NCL-mGFP to monitor its mobility within the nucleolus. This experiment revealed that 90 minutes of acetate removal, where we observed massive deacetylation (**Figure 2B-2D**), accelerated the half recovery times of NCL-mGFP (**Figure 5G-5H**). This result suggests that the mobility of NCL-mGFP within the nucelous was facilitated by acetyl-CoA depletion. Moreover, the precence of the class IIa inhibitor LMK235 but not the class I inhibitor Entinostat restored the half recovery times in acetate-depleted cells to smililar levels as in acetate-supplemented cells (**Figure 5G-5H**). We note that, when compared with LMK235, TMP195 exhibited only partial effects on the FRAP of NCL-mGFP. As also seen in the effect on acetylation of NCL (**Figure 5-figure supplement 2B**), this suggests that TMP195 might be less efficient in inhibiting the particular HDAC responsbile for deacetylation of NCL. Overall, these observations demonstrate that class IIa HDACs-mediated deacetylation alters the dynamics of nucleolar proteins, which are likely associated with changes in the phase state of the nucleolus.

### Acetylation-dependent regulation of the nucleolus in HCT116 cells

To extend our observations to additional cell types, we generated an HCT116 human colorectal carcinoma cell line-based ASA-KO cell line using the same CRISPR-mediated strategy as for HT1080 cells (**Figure 1-figure supplement 1A**). Again, viable *ACLY* KO cells were only achieved by supplementation of acetate in the culture media (**Figure 6-supplment figure 1A**). Acetate removal upregulated p53 in HCT116 ASA-KO but not ASA-WT cells and this was inhibited by the class IIa HDAC inhbitors (**Figure 6-supplment figure 1A-1B**). Our HCT116 cells harbor p53-YFP stably expressing at low levels under non stress conditions (Stewart-Ornstein and Lahav, 2017). Upon acetate withdrawal, p53-YFP expression was simillary induced as endogenous p53, adding further evidence for the post-transcriptional regulation of p53 upon acetyl-CoA depletion (**Figure 6-supplment figure 1A-1B**). Moreover, class IIa HDAC-dependent nucleolar remodelings were also obsereved in HCT116 ASA-KO cells (**Figure 6-supplment figure 1C**). These observations extend the generality of our findings regarding acetylation-mediated regulation of the nucleolar stress responses.

**Figure 6.**
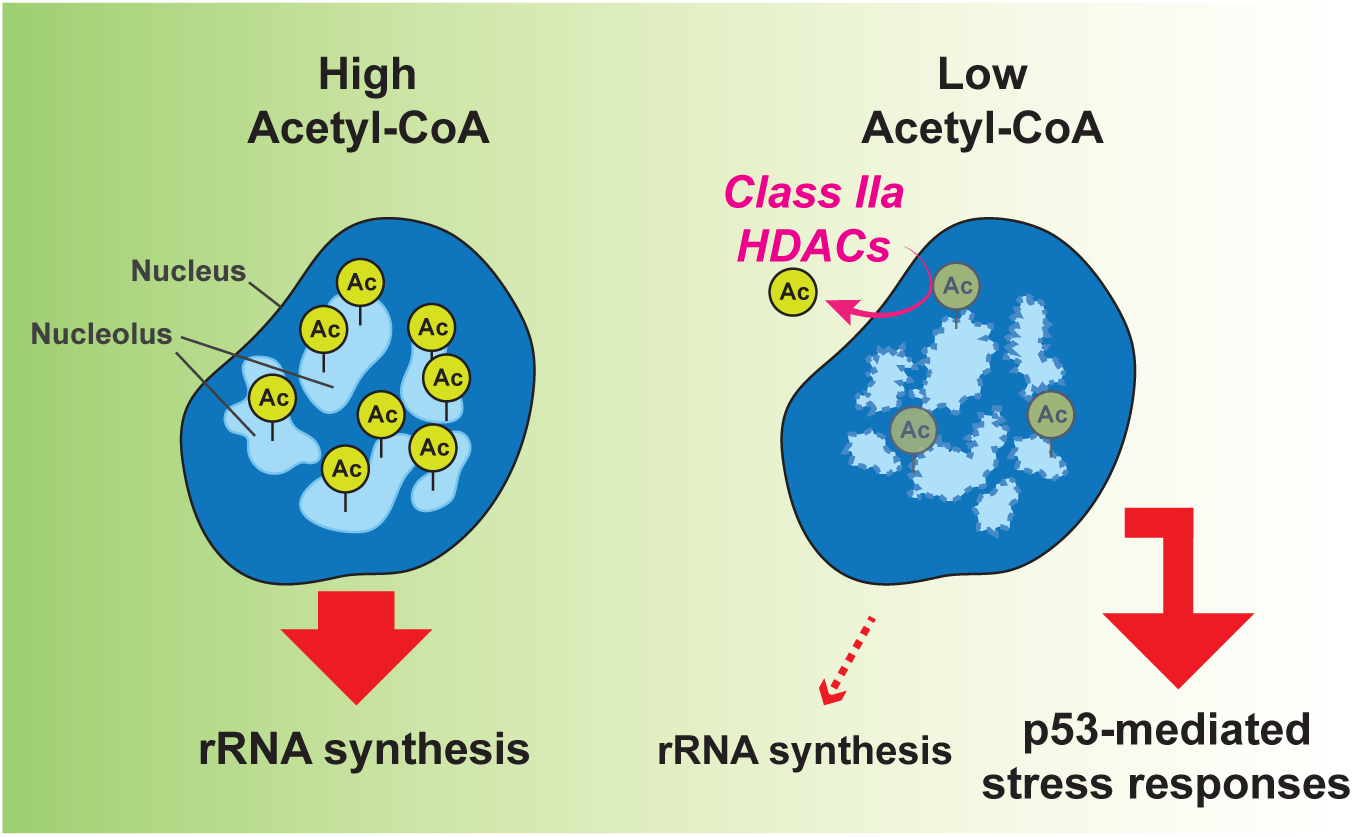
Model for the nucleolar responses upon nucleo-cytoplasmic acetyl-CoA fluctuations. See the main text for detail.

## Discussion

### The nucleolus as an acetyl-CoA responder

In this study, we demonstrated that the nucleolus rapidly responds to acetyl-CoA depletion. We propose a model for this cellular response to a decline in acetyl-CoA nucleo-cytoplasmic levels (**Figure 6**). In cells that contain high levels of acetyl-CoA, multiple nucleolar proteins are highly acetylated and these post-translational modifications play a crucial role in maintaining the nucleolar integrity, a phase separation state allowing for efficient rRNA synthesis. Once acetyl-CoA levels fall, class IIa HDACs deacetylate a significant number of nucleolar proteins. These reactions remodel the properties of the LLPS of the nucelolus, resulting in impairment of rRNA synthesis and induction of the p53-mediated stress responses. This model highlights an important role of the nucleolus as an organelle sensor for coupling acetyl-CoA fluctuations to cellular stress responses.

Accumulating evidence has illustrated that the nucleolus serves as a stress responsive organelle towards a variety of stresses including genotoxic stress, oxidative stress, heat shock, nutrient deprivations, virus infections and oncogene activations (Boulon et al., 2010; Deisenroth et al., 2016; Rubbi and Milner, 2003). These nucleolar stresses, as well as chemical or genetic interventions of ribosomal biogenesis, are known to induce morphological and functional alterations in the nucleolus, which are coupled with the activation of p53 or other signaling pathways in a stimulus- and cell context-dependent manner (Boulon et al., 2010; Chen and Stark, 2019). mRNA translation is known to be among the most energy demanding processes in mammalian cells and efficient ribosome biogenesis is nessesary for cell growth and proliferation (Buttgereit and Brand, 1995). Therefore, it is likely that a fall in acetyl-CoA levels signals a state of energetic stress and that cells acutely reduce ribosomal biogenesis in the nucleolus in order to limit energy consumption. Similar couplings of translational control and stress responses, but at different translational steps, are also observed in the integrated stress response (Pakos-Zebrucka et al., 2016).

This study uncovered that the metabolite acetyl-CoA can be a novel modulator of the nucleolus through protein acetylation. It has been reported that rRNA synthesis and rRNA processing are regulated by multiple mechanisms through neutrient signaling pathways to meet the cellular energy demands (Grummt, 2013). Toghether with these regulations, the cell likely employs this acetylation-mediated nucleolar stress response in order to help cope with physiological or pathlogical fluctuations in the cellular nutrient status.

### Nucleolus, acetylation and phase separation

Recent studies have revealed that PTMs can promote or oppose phase separation of biomolecular condensates through diverse mechanisms (Hofweber and Dormann, 2019; Snead and Gladfelter, 2019). Rapid and reversible alterations of PTMs that are incudced by various stimuli is capable of tuning the state of phase separation in a context-dependent manner. It has been recently reported that the LLPS-mediated formation of stress granules (SGs), membraneless organelles forming in response to stress, is modulated by acetylation of its component RNA helicase DDX3X (Saito et al., 2019). In this case, deacetylaton of DDX3X by HDAC6 is required for SGs maturation. Acetylation is also known to regulate the phase searation of pathogenic proteins which are involved in neurodegerative disorders, including Tau and TDP-43 (Carlomagno et al., 2017; Cohen et al., 2015; Ferreon et al., 2018). Hence, it is tempting to speculate that the acetylation status of the nucleolar components may also participate in regulating the nucleolar phase separation and thereby affect its functional state. We indeed observed enhanced mobility of GFP-NCL upon acetyl-CoA depletion (**Figure 5G-5H**), indicating alterations in the phase material property of the nucleolus, which is reported to be tightly linked to the nucleolar function (Zhu et al., 2019). In addition, our observations that acetyl-CoA depletion triggered redistribution of rRNA and NPM1, known contributors to the LLPS of the nucleolus, indicate changes in the phase state of the nucleolus (**Figure 3D-3F**) (Feric et al., 2016; Mitrea et al., 2016; Mitrea et al., 2018). Our acetylome analyses identified multiple acetylated lysine residues in various nucleolar protein (**Table S1 and Figure 5-supplment figure 1E**). The diversity of this list made it difficult to point out particular acetylation sites that are critical for the nucleolar integrity. It is likely that the entire balance of nucleolar protein acetylation may determine the propensity for LLPS. In this regard, we do not exclude a possible involvement of acetylation of non-nucleolar proteins that reside in the environment surrounding the nucleolus, including the nuclear lamina, actin cytoskeleton or chromatin, as these components are also known to influence the nucleolar structures and functions (Buchwalter and Hetzer, 2017; Feric and Brangwynne, 2013; Martin et al., 2009; Murayama et al., 2008; Sen Gupta and Sengupta, 2017).

### Class IIa HDACs regulate acetylation of nucleolar proteins

Compared with the class I HDACs or the class IIb HDAC6, substrates of the class IIa HDACs have been less characterized (Haberland et al., 2009; Parra, 2015). It has been previously shown that the class IIa HDACs have only minimal deacetylase activity on acetylated histones in vitro (Fischle et al., 2002; Lahm et al., 2007). Also, the transcriptional repressor activity by the class IIa HDACs expressed in cells or the enzymatic activity associated with immunoprecipitated class IIa HDACs are due to the deacetylase activity of the class I HDACs endogenously forming a complex with the class IIa HDACs (Fischle et al., 2002; Lahm et al., 2007). A catalytic tyrosine residue within the HDAC domain conserved in all other HDACs was found to be replaced with histidine in the class IIa HDACs, which can explain their weak catalytic activity towards canonical HDAC substrates (Lahm et al., 2007). Therefore, the importance of the catalytic activity of the class IIa HDACs for their biological functions has been questioned. Our HDAC inhibitor profiling however demonstrated that the acetyl-CoA depletion-dependent nucleolar alterations and p53 induction was selectively sensitive to class IIa HDAC inhibitors, indicating that a class IIa HDAC catalytic domain-dependent reactions exist in this context. Our acetylome analyses with TMP195 provide a list of potential substrates for class IIa HDACs (**Table S1**). This list highlights that various non-histone proteins including nucleolar proteins and ribosomal proteins are potential targets for this family of enzymes. Comparison with acetylome analyses using other class-selective HDAC inhibitors and in vitro reconstitution study will define bona fide substrates for the class IIa HDACs.

In conclusion, we demonstrated that our ASA-KO cell system can be a useful tool for analyzing acute responses to acetyl-CoA depletion. Our findings indicate that acetylation-mediated control of the nucleolar phase separation serves as an important hub linking acetyl-CoA metabolism to cellular stress responses. Furthermore, our findings highlight a potential application of the class IIa HDAC inhibitors for modulating nucleolar functions, which may provide novel insights for therapeutic interventions in a range of human diseases and in normal aging, where alterations or malfunctions of the nucleolus have been increasingly identified to play a role (Orsolic et al., 2016; Tiku and Antebi, 2018).

## Acknowledgments

We are grateful to members of Finkel lab for valuable discussion and technical supports and to the NHLBI Cores (Biochemistry Core, Bioinformatics and Computational Biology Core, DNA Sequencing and Genomics Core, Flow Cytometry Core, Light Microscopy Core, and Proteomics Core) and the Flow cytometry core at the McGawin Institute, University of Pittsburgh for experimental supports. We wish to acknowledge Duck-Yeon Lee at the NHLBI Biochemistry Core for HPLC instrumentation and calibration for acetyl-CoA and CoA. This work was supported by National Institutes of Health (Intramural Funds and 1 R01 HL142663 to T.F.), Foundation Leducq (to T.F.), and by the Aging Institute of the University of Pittsuburgh (to Y.S. and S.S.).

## Author Contributions

Y.S. and T.F. cocieved and led the project. Y.S., R.H. and S.S. performed experiments and analyzed data. H.K. contributed to developing the method for quantification of acetyl-CoA and CoA. S.G.W. conducted the lipid quantification and the data analysis. A.J.N. and M.P.S. conducted the acetylome experiments and the data analysis. Y.L. supervised the RNA sequensing experiments. F.S. and M.P. computationally analyzed the RNA sequensing data. G.W. and M.G. conducted the TMT proteomics and the data analysis. D.A.M. conducted the microscopy image data analysis. S.C.W supervised and M.J.C. conducted the FRAP experiments. J.S.-O. contributed to developing the HCT116 p53-YFP cell system. Y.S. and S.S. visualized data and schemes. Y.S. wrote the initial draft with input of other authors. T.F. and all other authors reviewed and editied the manuscript.

## Materials and Methods

### Cell culture

HT1080 human fibrosarcoma cells (ATCC, CCL-121) were cultured at 37°C with 5% CO_2_ in Minimum Essential Medium Eagle (with Earle’s salts, L-glutamine, and sodium bicarbonate) (Sigma-Aldrich, St. Louis, MO), supplemented with 10% heat inactivated Fetal Bovine Serum (Thermo Fisher Scientific, Asheville, NC, Gibco or VWR Life Science, Randor, PA), 1x Penicillin Streptomycin Solution (Corning Incorporated, Corning, NY) and 1x MEM Non-essential Amino Acid Solution (Sigma). Sodium Acetate (Sigma) was supplemented in the culture medium for the *ACLY* sgRNA-targeted cells. Dialyzed FBS was purchased from Thermo Fisher Scientific (Gibco). 293T cells (Lenti-X, Takara, San Francisco, CA) and HCT116 cells (Stewart-Ornstein and Lahav, 2017)were cultured in the same condition as HT1080 cells but in Dulbeccos’s Modified Eagle Medium (Gibco) supplemented with 10% FBS, 1 mM Sodium pyruvate (Gibco), non-essential amino acids (Gibco) and GlutaMAX (Gibco).

### Plasmid constructions and lentivirus productions

Oligo duplexes containing a single guide (sg)RNA sequence for ACLY (as shown in Figure 1-figure supplement 1A) were inserted into pCas9(BB)-2A-GFP (a gift from Feng Zhang, Addgene #48138) following the published procedures (Ran et al., 2013) to generate genome-edited cells using the CRISPR-Cas9 system. For lentivirus-mediated gene expression, a coding sequence (CDS) of human ACLY was PCR amplified from a HeLa cDNA pool and inserted into the *EcoR*I-digested pLVX-puro vector (Takara) using the In-Fusion cloning system (Takara). A CDS of RPL11 was amplified from an HT1080 cDNA pool and inserted into the *EcoR*I/*Bgl*II-digested p3xFLAG-CMV-7.1 (Sigma) to attach the 3xFLAG tag at its N-terminus. The 3xFLAG-RPL11 sequence was then inserted into the *EcoR*I-digested pLVX-puro vector. The NCL CDS was amplified from the GFP-Nucleolin plasmid (a gift form Michael Kastan, Addgene #28176)(Takagi et al., 2005). The NCL-3xFLAG fragment was amplified using primers that attach the 3xFLAG at NCL’s C-terminus and was inserted into the *EcoR*I-digested pLVX-puro vector. For the NCL-mGFP expressing constract, the NCL-CDS was amplified from the GFP-Nucleolin plasmid and monomeric GFP (mGFP) was amplified from the LAMP1-mGFP plasmid (a gift from Esteban Dell’Angelica, Addgene #34831)(Falcon-Perez et al., 2005) and these fragments were inserted into the *EcoR*I-digested pLVX-puro vector. Lentivirus was produced in 293T cells by transfecting the pLVX-puro vectors, psPAX2 (a gift from Didier Trono, Addgene #12260) and pCMV-VSV-G (a gift from Bob Weinberg, Addgene #8454) (Stewart et al., 2003).

### Generation of the *ACLY* genome-edited cell lines by CRISPR-Cas9

The pCas9(BB)-2A-GFP plasmids containing the sgRNA sequence targeting *ACLY* were transfected into HT1080 cells or HCT116 p53-YFP cells using Lipofectamine LTX (Thermo Fisher Scientific) or X-treamGENE (Sigma) according to the manufacturer’s instruction. The next day, GFP positive cells were collected using the FACS Fusion sorter (BD Biosciences) and cultured in three different medium conditons; standard culture media or the same media with 2 or 20 mM sodium acetate as illustrated in Figure 1-figure supplement 1A. Cell clones expanded from each condition were screened by genotyping PCR (the forward primer; gtggctgaagagctatgtccag and the reverse primer; cctctgcttgtgcacatctgtc) combined with *Xho*I digestion (as illustrated in Figure 1-figure supplement 1B) and by immunoblotting.

### Immunoblotting

ASA cells were plated at a density of 8 × 10^4^ cells per well in a 6 well plate and cultured in the 20 mM sodium acetate-supplemented culture medium (10% FBS). The next day, the cells were washed with PBS twice and 1% FBS culture medium once, and then cultured in the media containing 10% or 1% FBS with or without 20 mM sodium acetate, or with indicated chemicals according to experiments. After indicated time periods, the cells were washed with PBS or 1%FBS culture medium once and lysed in 150 ul of 1x NuPAGE LDS sample buffer (Thermo Fisher Scientific) supplemented with 10 mM DTT. The samples were shaken at max speed in a table top incubator at room temperature for 10 min, boiled at 98°C for 5 min, and centrifuged at 15,000 rpm for 5 min. The supernatant containing approximately 15-30 μg protein was separated by SDS-PAGE using a 4-20% gradient Mini-PROTEAN TGX Precast Gel (Bio-Rad, Hercules, CA) and then transferred to a 0.2 μM nitrocellulose membrane. The membrane was blocked with Odyssey Blocking Buffer (LI-COR, Lincoln, NE) and incubated with the indicated primary antibodies at 4°C overnight. After washing with PBS-T (PBS + 0.05% Tween-20), the membrane was incubated with near-infrared fluorescent IRDye secondary antibodies (LI-COR) or HRP-conjugated secondary antibodies (Thermo Fisher Scientific) and washed again with PBS-T. Detection was performed with either the ODYSSEY CLx Infrared Imaging System together with Image Studio Lite version 5.2 Software (LI-COR), or iBright CL1000 Imaging System (Thermo Fisher Scientific). For immunoblotting of BOP1 and RRP1, the cells were lysed with 1% Triton buffer [1% Triton-X100, 150 mM NaCl, 50 mM Tris-HCl pH 7.4, 1 mM EDTA, Phosphatase inhibitors (PhosSTOP, Sigma) and protease inhibitors (cOmplete, Sigma)]. After centrifugation, the supernatant was mixed with 4x NuPAGE LDS sample buffer and boiled. For siRNA-mediated knockdown experiments, silencer siRNAs (5 nM) for indicated genes or Negative Control siRNA #1 (Ambion, AM4611) were transfected with Lipofectamin RNAi MAX transfection reagent (Thermo Fisher Scientific, 13778075) into 4 x10^4^ cells in a 6 well plate. The next day, the culture media were replaced with fresh media, and the day after that the cells were subjected to immunoblotting experiments. Following Silencer Select Pre-designed siRNAs were used; RPL5 (ID s56731), RPL11 (ID s12168).

### Immunoprecipitation

ASA-KO17 cells and the ASA-KO17 cells stably expressing 3xFLAG-RPL11 were plated at a density of 7 × 10^5^ cells per 10 cm culture dish and cultured in the 20 mM sodium acetate-supplemented culture medium (10% FBS). The next day, the cells were washed with PBS twice and 1% FBS culture medium once, and then cultured in 1% FBS culture medium with or without 20 mM sodium acetate (two 10 cm dishes were prepared for each condition). After indicated time periods, the cells were washed with PBS and lysed in 500 μl of 1% Triton buffer [1% Triton-X100, 150 mM NaCl, 50 mM Tris-HCl pH 7.4, 1 mM EDTA, Phosphatase inhibitors (PhosSTOP, Sigma) and protease inhibitors (cOmplete, Sigma)]. After centrifugation, 30 μl of the supernatant was taken for “Lyasate” sample and the rest of the supernatant was mixed with 10 μl of Anti-FLAG M2 Magnetic beads (Sigma). After 30 min of incubation at 4°C, the beads were washed with 1% Trition buffer three times and then incubated in 30 μl of 1% Trition buffer with 0.2 μg/μl of 3xFLAG peptide (Sigma) at room temperature for 20 min. Then the supernatant was mixed with 4x NuPAGE LDS sample buffer (Thermo Fisher Scientific) supplemented with 10 mM DTT and boiled. The immunoprecipitation assay was performed using an acetyl-Lysine motif antibody to detect acetylated protein. The ASA-KO17 cells stably expressing 3xFLAG-RPL11 or NCL-3xFLAG were plated, stimulated, and lysed as the FLAG immunoprecipitation assay described above. The supernatant was mixed with either 0.8 μg Rabbit IgG (Santa Cruz, Dallas, TX, sc-2027) or 4 μl of the acetyl-lysine motif antibody (Cell Signaling Technology, Danvers, MA, #9814) and incubated over night at 4°C. Then the supernatant was mixed with 10 μl of Protein A/G Magnetic Beads (Thermo Fisher Scientific) and incubated for 1 hour at 4°C. The beads were washed with 1% Trition buffer three times and mixed with 35 μl of 1x NuPAGE LDS sample buffer supplemented with 10 mM DTT and shaken at room temperature for 10 min. Then the supernatant was subjected to SDS-PAGE.

### Cell viability analysis (WST-1 assay)

Cells were plated at a density of 5,000 cells per well in a 6 well plate and cultured in the culture medium (10% FBS) with or without 20 mM sodium acetate. The next day, the cells were washed with PBS twice and 1% FBS culture medium once, and then cultured in the media containing 10% FBS, 1% FBS or 10% dialyzed FBS with or without 20 mM sodium acetate. 24 hours later, the culture media were replaced with 10% FBS culture medium with or without 20 mM sodium acetate and the cells were cultured for another 2 days. Then, the media were replaced with 10% FBS medium containing 50 µM WST-1 (Dojindo, Gaithersburg, MD) and 20 µM 1-methoxy PMS (Dojindo), and the cells were incubated for 2 hours at 37°C in the cell culture incubator before the absorbance at 440 nm in the culture media (100 μl aliquots transferred in a 96 well plate) was measured using Spectra Max Plus 384 microplate Reader (Molecular Devices, Sunnyvale, CA).

### Quantification of Acetyl-CoA and Coenzyme A in cell lysate by HPLC

ASA-WT5 and ASA-KO17 cells were plated at a density of 4 × 10^5^ cells per 10 cm culture dish and cultured in the 20 mM sodium acetate-supplemented culture medium (10% FBS) for 1 day (10 dishes were prepared for each cell line). The cells were then washed with PBS twice and cultured in 1 % FBS culture media with or without 20 mM sodium acetate (5 dishes were prepared for each condition). After 4 hours, cells were washed with PBS, detached by trypsin, and resuspended in the culture media (approximately 1.5-2.0 × 10^6^ cells). Cell pellet was collected by centrifugation, washed with PBS twice, and resuspended in 100 μl of 5% 5-Sulfosalicylic acid (Sigma) solution. To permeabilize the cells, samples were frozen in liquid nitrogen and thawed on ice. This freeze-thaw cycle was repeated twice. After centrifugation, the supernatant was filtered using Ultrafree-MC LH Centrifugal Filter (Millipore, Billerica, MA, UFC30LH25). Then, the samples were transferred into SUN-SRi Glass Microsampling Vials (Thermo Fisher Scientific, 14-823-359) with SUN-SRi 11mm Snap Caps (Thermo Fisher Scientific, 14-823-379) and 80 μl of each sample was separated using an Agilent 1100 HPLC (Agilent Technologies, Wilmington, DE) equipped with a reverse phase column, Luna 3 μm C18(2) 100 Å, 50 × 4.6mm, 3 μm (Phenomenex, Los Angeles, CA). The HPLC-reverse phase column was calibrated with acetyl-CoA (Sigma, A2056) and coenzyme A (Sigma, C3144). The mobile phase was described previously (Shibata et al., 2012) and consisted of two eluents: 75 mM sodium acetate and 100 mM NaH_2_PO_4_ pH 4.6 (buffer A). Methanol was added to the buffer A at a ratio of 70:30 (v/v) = buffer A:methanol (buffer B). The gradient started with 10% of buffer B and was run under following conditions; 10 min at up to 40% of buffer B, 13 min at up to 68% of buffer B, 23 min at up to 72% of buffer B, 28 min at up to 100% of buffer B, and held for an additional 5 min. The initial condition was restored after 10 min with 10% of buffer B. The flow rate was 0.6 ml/min and the detection was performed at 259 nm. UV chromatograms were analyzed using the software ChemStation version A.10.01 (Agilent Technologies).

### Quantification of Cholesterol and fatty acids

ASA-KO17 cells were plated and stimulated and the cell pellet (approximately 1.1-2.8 × 10^6^ cells) was collected as described in the section for quantification of acetyl-CoA. For measurement of cholesterol and fatty acids, 500 µL of aqueous cell lysate was extracted with 2 mL of chloroform:methanol (2:1) after the addition of 10 µL of a deuterated fatty acid internal standard mix (50 ppm) and 20 µL of cholesterol-d_7_ (10 ppm). The samples were vortexed and centrifuged at 2500 rpm for 10 min at 4°C and the organic layer was transferred into two new glass vials in equal volumes for fatty acid derivatization and cholesterol analysis. Both vials were dried under N_2_. Samples for cholesterol analysis were reconstituted in 100 µL of chloroform:methanol (2:1) and 10 µL was injected into a Shimadzu LC with CTC PAL autosampler coupled to a Sciex 5000 triple quadrupole mass analyzer. Analytes were separated on a C18(2) Luna column (2.1 × 100 mm, Phenomenex) at a flow rate of 0.63 mL/min using 50:50 H_2_O:ACN with 0.1% formic acid for solvent A and 40:60 ACN:IPA with 0.1% formic acid for solvent B. The gradient started at 50% B and increased to 100% B over 6 minutes before returning to initial conditions for column equilibration. Cholesterol and cholesterol-d_7_ were detected by selective reaction monitoring using 369→147 and 376→147 transitions, respectively. Peak area ratios of cholesterol to cholesterol-d_7_ were normalized to cell number and are reported as relative amount.

Fatty acids were derivatized for LC-MS analysis using a Vanquish UPLC coupled to a Q Exactive mass spectrometer (Thermo Fisher Scientific). Derivatization of the samples was conducted as described (Li and Franke, 2011). To each sample, 200 µL of oxalyl chloride (2 M in dichloromethane) was added and the mixture was incubated for 5 min at 65°C. Samples were dried under N_2_ and 150 µL of 3-picolylamine (1% v/v in ACN) was added for a 5 min incubation at room temperature to form the 3-picolylamide (PA) ester. Samples were dried a final time under N_2_ and reconstituted in 1 mL MeOH and 5 µL was injected onto the Vanquish-QE system. Fatty acid-PA esters were separated on a Phenomenex C8 column (2.1 × 150 mm, 5 µ pore size) using H_2_O + 0.1% acetic acid for solvent A and ACN + 0.1% acetic acid for solvent B. The gradient started at 65% B and increased linearly to 85% B at 10 min and was held for 1 min before ramping to 100% B for 2min. Finally, the gradient returned to 65% B for a 2 min equilibration. Samples were analyzed using full scan accurate mass at a resolution of 70K. Peak area ratios of fatty acid-PA ester to corresponding internal standard were normalized to cell number and reported as relative amount.

### Acetylome analysis

ASA-KO17 cells were plated at a density of 6 × 10^6^ cells per 15 cm culture dish and cultured in the 20 mM sodium acetate-supplemented culture media (10% FBS) for 1 day. The cells were then washed with PBS twice and 1% FBS culture medium once, and cultured in 1% FBS culture media with or without 20 mM sodium acetate, or without acetate but with 50 μM TMP195 (21 dishes were prepared for each condition). After 90 min, the cells were washed with 10 ml of cold PBS once and lysed in 10 ml of PTMScan Urea Lysis Buffer [20 mM HEPES (pH 8.0), 9.0 M urea, 1 mM sodium orthovanadate (activated), 2.5 mM sodium pyrophosphate, 1 mM *β*-glycerol-phosphate] (Cell Signaling Technology). Samples were frozen in dry ice/ethanol and then analyzed using the PTMScan method (Cell Signaling Technology) as previously described (Rush et al., 2005; Stokes et al., 2015). Briefly, the lysates were sonicated, centrifuged, reduced with DTT, and alkylated with iodoacetamide. 15 mg of total protein for each sample was digested with trypsin and purified over C18 columns for enrichment with the Acetyl-Lysine Motif Antibody (#13416). Enriched peptides were purified over C18 STAGE tips. Enriched peptides were subjected to secondary digest with trypsin and second STAGE tip prior to LC-MS/MS analysis. Replicate injections of each sample were run non-sequentially on the instrument. Peptides were eluted using a 90-minute linear gradient of acetonitrile in 0.125% formic acid delivered at 280 nL/min. Tandem mass spectra were collected in a data-dependent manner with a Thermo Orbitrap Fusion™ Lumos™ Tribrid™ mass spectrometer using a top-twenty MS/MS method, a dynamic repeat count of one, and a repeat duration of 30 sec. Real time recalibration of mass error was performed using lock mass with a singly charged polysiloxane ion m/z = 371.101237. MS/MS spectra were evaluated using SEQUEST and the Core platform from Harvard University. Files were searched against the SwissProt *Homo sapiens* FASTA database. A mass accuracy of +/-5 ppm was used for precursor ions and 0.02 Da for product ions. Enzyme specificity was limited to trypsin, with at least one tryptic (K- or R-containing) terminus required per peptide and up to four mis-cleavages allowed. Cysteine carboxamidomethylation was specified as a static modification, oxidation of methionine and acetylation on lysine residues were allowed as variable modifications. Reverse decoy databases were included for all searches to estimate false discovery rates, and filtered using a 2.5% FDR in the Linear Discriminant module of Core. Peptides were also manually filtered using a −/+ 5ppm mass error range and presence of an acetylated lysine residue. All quantitative results were generated using Skyline to extract the integrated peak area of the corresponding peptide assignments. Accuracy of quantitative data was ensured by manual review in Skyline or in the ion chromatogram files. GO analysis was done using fuctional annotation in DAVID Bioinformatics Resources (david.ncifcrf.gov).

### RNA sequencing analysis

ASA-KO17 cells were plated at a density of 4 × 10^5^ cells per 10 cm culture dish and cultured in the 20 mM sodium acetate-supplemented FBS culture media (10% FBS) for 1 day. The cells were then washed with PBS twice, and cultured in the culture media containing 10% or 1% FBS with or without 20 mM sodium acetate (2 dishes were prepared for each condition). After 4 hours, cells were washed with PBS, collected in 1 mM EDTA in PBS and total RNAs were isolated using RNeasy Mini Kit (Qiagen, 74106) according to the manufacturer’s instruction. The sequencing libraries were constructed from 100 ng – 500 ng of total RNA using the Illumina’s TruSeq Stranded Total RNA kit with Ribo-Zero following the manufacturer’s instruction. The fragment size of RNAseq libraries was verified using the Agilent 2100 Bioanalyzer (Agilent) and the concentrations were determined using Qubit instrument (LifeTech). The libraries were loaded onto the Illumina HiSeq 3000 for 2×75 bp paired end read sequencing. The fastq files were generated using the bcl2fastq software for further data analysis. Data analysis: After quality assessment, adapter and low-quality bases trimming, reads were aligned to the reference genome using the latest version of HISAT2 (Kim et al., 2015), which sequentially aligns reads to the known transcriptome and genome using the splice-aware aligner built upon HISAT2. Only uniquely mapped paired-end reads were then used for subsequent analyses. FeatureCounts (Liao et al., 2014) was used for gene level abundance estimation. Differential expression analysis were then carried out using open source Limma R package (Ritchie et al., 2015). Limma-voom was employed to implement a gene-wise linear modelling which processes the read counts into log_2_ counts per million (logCPM) with associated precision weights. The logCPM values were normalized between samples using trimmed mean of M-values (TMM). We adjust for multiple testing by reporting the FDR q-values for each feature. Features with q < 5% were declared as genome-wide significant.. GO analysis was done using fuctional annotation in DAVID Bioinformatics Resources (david.ncifcrf.gov).

### Tandem mass spectromety analysis

ASA-KO17 cells were plated at a density of 4 × 10^5^ cells per 10 cm culture dish and cultured in the 20 mM sodium acetate-supplemented culture media (10% FBS) for 1 day. The cells were then washed with PBS twice, and cultured in 1% FBS culture media with or without 20 mM sodium acetate (2 or 3 dishes were prepared for each condition). After 4 hours, cells were washed with ice-cold PBS three times, harvested with 1 mM EDTA in PBS, collected by centrifugation, and lysed with RIPA buffer [1% NP-40, 150 mM NaCl, 50 mM Tris-HCl pH 7.4, 0.1% SDS, 0.5% Na-DOC, Phosphatase inhibitor (PhosSTOP, Sigma) and protease inhibitor (cOmplete, Sigma)]. After centrifugation, the supernatant containing 100 μg protein was used for the tandem mass spectrometry analysis. The samples were reduced, alkylated, digested, and labeled according to instructions for the TMT 10plex kit (Thermo Fisher Scientific). TMT-labeled peptides were mixed, desalted, and fractionated on an Agilent 1200 HPLC system to 24 fractions using basic reversed-phase chromatography. LC-MS/MS was performed on a Dionex UltiMate 3000 rapid separation nano UHPLC system (Thermo Fisher Scientific) coupled online to an Orbitrap Fusion Lumos tribrid mass spectrometer (Thermo Fisher Scientific). Peptides were first loaded onto a nano trap column (Acclaim PepMap100 C18, 3 μm, 100Å, 75 μm i.d. × 2 cm, Thermo Fisher Scientific), and then separated on a reversed-phase EASY-Spray analytical column (PepMap RSLC C18, 2 μm, 75 μm i.d. × 50 cm, Thermo Fisher Scientific) using a linear gradient of 4-32% B (buffer A: 0.1% formic acid in water; buffer B: 0.1% formic acid in acetonitrile) for 100 min. The mass spectrometer was equipped with a nano EASY-Spray ionization source, and eluted peptides were brought into gas-phase ions by electrospray ionization and analyzed in the orbitrap. High-resolution survey MS scans and HCD fragment MS/MS spectra were acquired in a data-dependent manner with a cycle time of 3 s. Dynamic exclusion was enabled. Raw data files generated from LC-MS/MS were analyzed using a Proteome Discoverer v2.2 software package (Thermo Fisher Scientific) and the Mascot search engine (Matrix Science). The following database search criteria were set to: database, SwissProt human; enzyme, trypsin; max miscleavages, 2; variable modifications, oxidation (M), deamidation (NQ); fixed modifications, TMT (K, N-term), carbamidomethylation (C); peptide precursor mass tolerance, 20 ppm; MS/MS fragment mass tolerance, 0.03 Da. Peptide-spectrum matches (PSMs) were filtered to achieve an estimated false discovery rate (FDR) of 1% based on a target-decoy database search strategy. The relative abundance of a protein or peptide in different samples was estimated by TMT reporter ion intensities.

### Light Microscopy analysis

Cells were plated at a density of 4 × 10^4^ cells per well in a 4 well chamber slide (Nunc Lab-Tek II Chamber slide system, Thermo Fisher Scientific), and cultured in the 20 mM sodium acetate-supplemented culture medium (10% FBS). The next day, the cells were washed with PBS twice and 1% FBS culture medium once, and cultured in 1% FBS culture media with or without 20 mM sodium acetate, or with or without indicated compounds. After indicated periods, the cells were washed with PBS, fixed with 4% formaldehyde/PBS for 10 min, permeabilized with 1% Triton-X/PBS for 15 min and blocked with 2% BSA/PBS for 30 min at room temperature. Primary antibodies in 2% BSA/PBS were incubated overnight at 4°C and secondary antibodies (Thermo Fisher Scientific, Alexa Fluor) were incubated for 2 hours at room temperature and washed with PBS-T. 1 μM Nucleolus bright red (Dojindo) was treated in PBS for 10 min and washed with PBS just before the observation. For FUrd incorporation assay, 5-Fluorouridine (1 mM) (Sigma) was added in the culture media 15 min before fixation. The FUrd signal was detected by staining with anti-BrdU [BU1/75 (ICR1)] Rat mAb (abcam, Boston, MA, ab6326) and Alexa Fluor 488 anti-Rat IgG (Jackson Labratory, Bar Harbor, ME, 712-547-003). Fluorescent images were acquired using Leica SP8 LIGHTNING Confocal Microscope (Leica, Buffalo Grove, IL) equipped with a 63 × /1.4 NA oil immersion objective and driven by LAS X software. Image quantifications were performed with LAS X software. Fluorescent images in Figure 4S were acquired using a Zeiss LSM 780 Confocal Microscope (Carl Zeiss, Dublin, CA) equipped with a 63 × /1.4 NA oil immersion objective and driven by ZEN software.

### FRAP analysis

ASA-KO17 cells were plated at a density of 2 × 10^5^ cells per well in a 35 mm glass bottom dish (No. 1.5 Coverslip, 14 mm glass diameter, uncoated, MatTek), and cultured in the 20 mM sodium acetate-supplemented culture medium (10% FBS). The next day, the cells were washed with PBS twice and 1% FBS culture medium once, and cultured in 1% FBS culture media with or without 20 mM sodium acetate, or with or without indicated compounds. 90 minutes later, confocal images were taken on a Leica SP8 confocal microscope with a HC PL APO 1.30 NA 93x glycerine objective using a Acousto-Optic Beam Splitter (AOBS) emission system on LASX software (Leica, v3.5.2.18963). Images were acquired at 1.54 frames per second, with 132nm pixels using a galvo scanner at 400 Hz. Marker mobility was measured using the FRAP module, bleaching within the ROI with three passes of high power 405nm laser. Immediately post-bleach, samples were imaged continuously for approximately 50 sec. T1/2 was measured using Elements (Nikon, Melville, NY, v5.210) time measurement tool.

### Reagents and antibodies

Reagents were sourced as indicated: Tricostatin A (Sigma, T1952), Vorinostat (SAHA, MK0683) (Selleck Chemicals, Houston, TX, S1047), Entinostat (MS-275) (Selleck Chemicals, S1053), Mocetinostat (Cayman Chemical, Ann Arbor, MI, 18287), RGFP966 (Cayman Chemical, 16917), TMP195 (Cayman Chemical, 23242), LMK235 (Cayman Chemical, 14969), Tubastatin A (Apex Biomedical, Clackamas, OR, A4101), Camptothecin (Selleck Chemicals, S1288), Actinomycin D (Sigma, A1410).

Antibodies were sourced as indicated:

ATP-Citrate Lyase Rabbit polyAb (1:1000, Cell Signaling Technology, #4332)

Histone H3 (D1H2) XP^®^ Rabbit mAb (1:1000, Cell Signaling Technology, #4499)

Acetyl-Histone H3 (Lys9) (C5B11) Rabbit mAb (1:1000, Cell Signaling Technology, #9649)

Acetyl-Histone H3 (Lys27) (D5E4) XP^®^ Rabbit mAb (1:1000, Cell Signaling Technology, #8173)

Acetyl-Histone H4 (Lys8) Rabbit polyAb (1:1000, Cell Signaling Technology, #2594)

Phospho-Histone H2A.X (Ser139) (20E3) Rabbit mAb (1:1000 or 1:200, Cell Signaling Technology, #9718)

Histone H2A (D6O3A) Rabbit mAb (1:1000, Cell Signaling Technology, #12349)

Acetylated-Lysine Antibody Rabbit polyAb (1:1000, Cell Signaling Technology, #9814)

α-Tubulin (clone DM1A) Mouse mAb (1:5000, Sigma, T6199)

Acetyl-α-Tubulin (Lys40) (D20G3) XP^®^ Rabbit mAb (1:1000, Cell Signaling Technology, #5335)

p53 (DO-1) Mouse mAb (1:5000, Santa Cruz, sc-126)

GLTSCR2/PICT1 Rabbit polyAb (1:1000, Cell Signaling Technology,73225)

MDM2 (SMP14) Mouse mAb (1:1000, Santa Cruz, sc-965)

RPL11 (D1P5N) Rabbit polyAb (1:1000, Cell Signaling Technology,18163)

RPL5 Rabbit polyAb (1:1000, Cell Signaling Technology,14568)

anti-BrdU [BU1/75 (ICR1)] Rat mAb (1:500, abcam, ab6326)

BOP1 (E-1) Mouse mAb (1:250, Santa Cruz, sc-390672)

RRP1 Rabbit polyAb (1:200, GeneTex, Irvine, CA, GTX115107)

UBF (F-9) Mouse mAb (1:500, Santa Cruz, sc-13125)

FBL (G-8) Mouse mAb (1:500, Santa Cruz, sc-374022)

FBL (B-1) Mouse mAb (1:500, Santa Cruz, sc-166001)

NCL/C23 (H-6) Mouse mAb (1:500, Santa Cruz, sc-55486)

NCL (D4C7O) Rabbit mAb (1:1000, Cell Signaling Technology, #14574)

NPM1 Rabbit polyAb (1:200,, Cell Signaling Technology, #3542)

Anti-FLAG M2 Mouse mAb (1:1000, Sigma, F3165)

---

**Figure 1-figure supplement 1.**
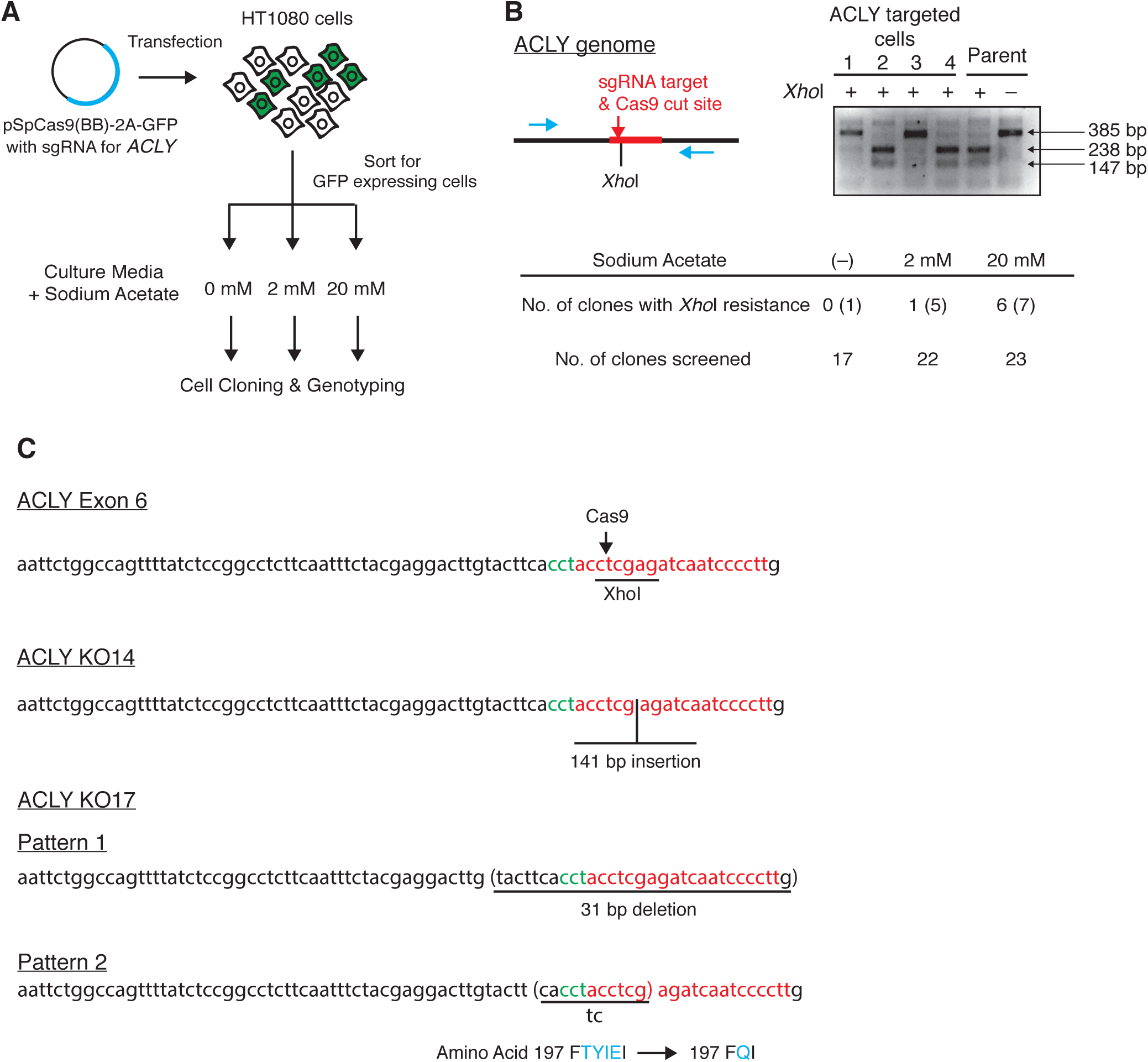
(A) Schema of genome editing for *ACLY*. A GFP-tagged sgRNA/Cas9 plasmid targeting *ACLY* was transfected into HT1080 cells. Following cell sorting by flow cytometry for GFP positive transfected cells, cells were recovered in either standard medium (no added acetate) or 2 or 20 mM sodium acetate-supplemented media. Recovered cell clones were genotyped to identify genome editing for *ACLY* as in “B”. (B) Schema of genotyping for *ACLY* genome-edited cell clones and representative genotyping results (top left). The sgRNA-targeted and Cas9 cleavage site in *ACLY* exon 6, which contains a single *Xho*I site overlapped with a Cas9 cleavage site, was PCR amplified by flanked primers (blue). *Xho*I digestion of the PCR product (385 bp) provided 238 bp and 147 bp fragments when the *Xho*I site was intact (top right). *Xho*I resistance was indicative of insertion or deletion (in/del) by CRISPR-mediated non-homologous end joining (NHEJ). Numbers of clones which exhibited complete (or partial) *Xho*I resistance and numbers of clones screened were shown for each culture media condition (bottom). (C) The sequence of exon 6 in wildtype *ACLY* and the edited sequences in ASA-KO14 and KO17 cells. The sgRNA target site (red), PAM motif (green), Cas9 cut site (arrow head), *Xho*I site, and detected in/del were indicated. We detected only a single pattern of insertion in ASA-KO14 cells. Note that one allele of ASA-KO17 possesses an indel which causes a frame shift mutation replacing amino acids “T-Y-I-E” with “Q”. This might explain the detected ACLY peptides in the TMT proteomics using ASA-KO17 cells (Table S3).

**Figure 2-figure supplement 1.**
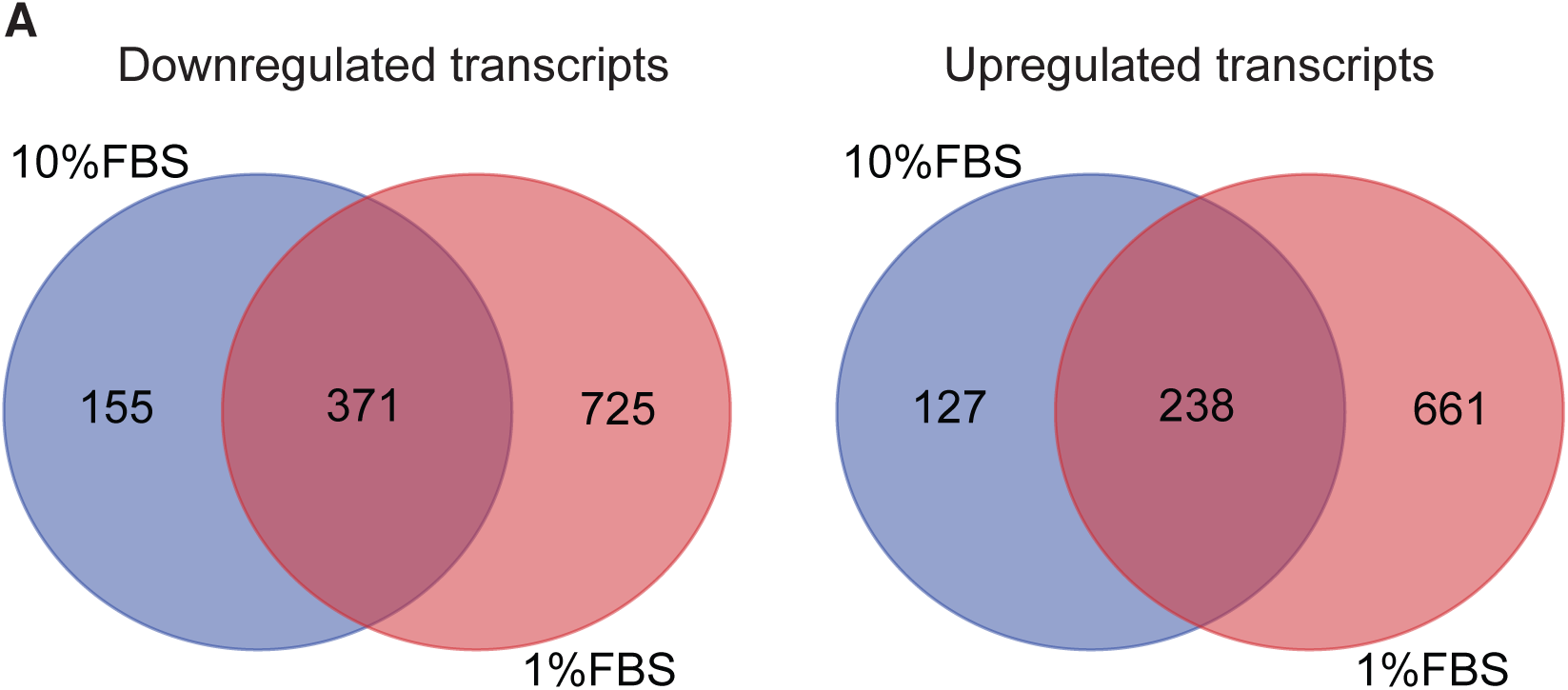
(A) Venn diagram for downregulated transcripts (log_2_ FC [(-) Acetate / (+) Acetate] < −1.0. q<0.05) and upregulated transcripts (log_2_ FC [(-) Acetate / (+) Acetate] >-1.0. q<0.05) in the RNA sequencing with the 10% and 1% FBS conditions.

**Figure 3-figure supplement 1.**
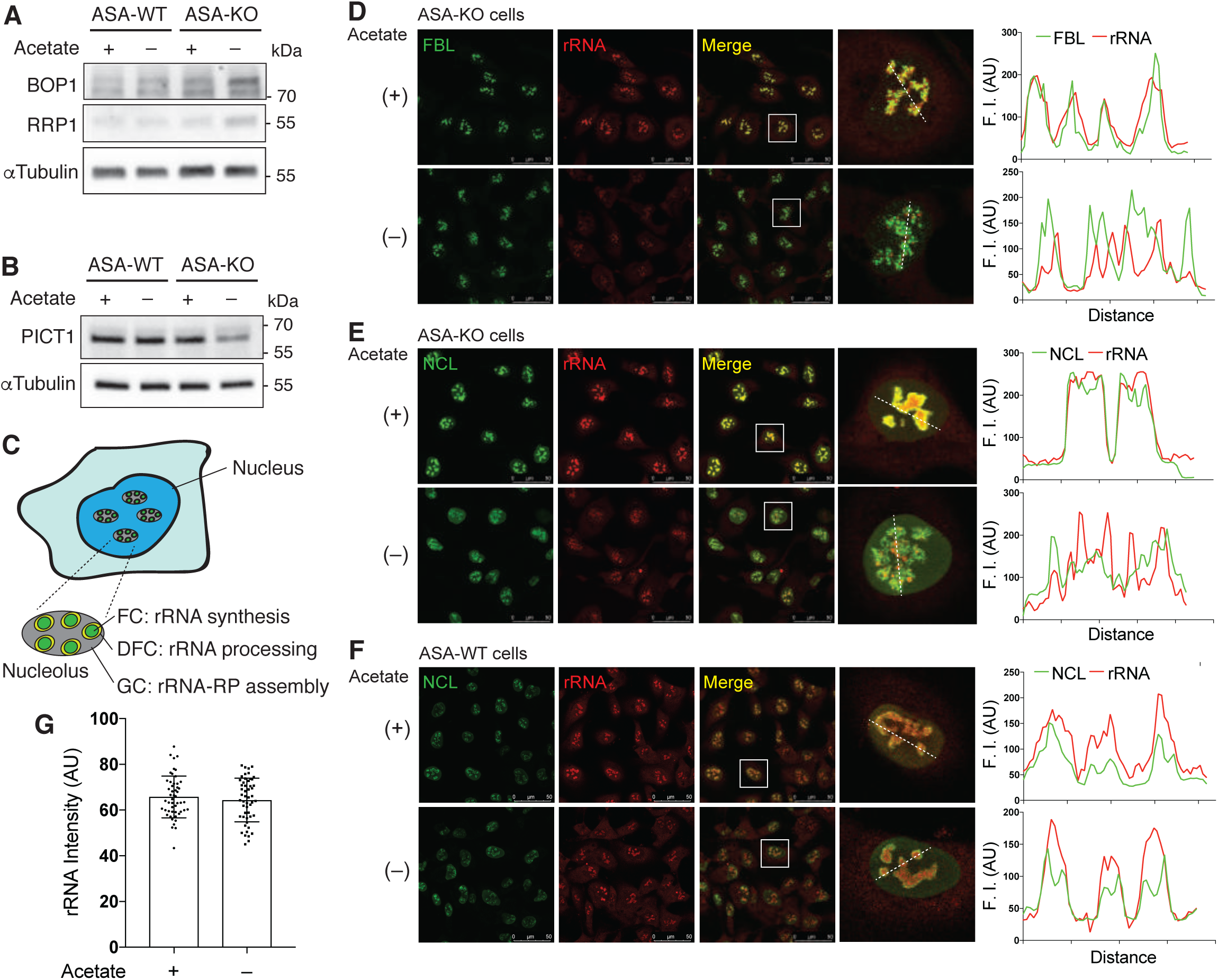
(A-B) Immunoblotting for nucleolar protein BOP1 and RRP1 (A) and PICT1 (B) in ASA-WT and ASA-KO cells cultured in 10% or 1% FBS containing media with or without acetate for 4 hours. *α*Tubulin was used as a loading control. (C) Schematic representation of the nucleolus consisting of fibrillar center (FC; green), dense fibrillar component (DFC; yellow) and granular component (GC; gray). (D-F) Immunostaining for FBL (D) and NCL (E and F) along with rRNA staining in ASA-KO (E and E) and ASK-WT (F) cells cultured in 1% FBS containing media with or without acetate for 4 hours. Magnified nuclear images (surrounded by a white square) are shown. Line profiles for indicated fluorescent intensities (F. I) determined along the white dashed lines are shown to the right. AU, arbitrary unit. (G) Quantification of the mean fluorescent intensity of the rRNA singal per nucleus in (F). Data are shown as mean ± S.D. (n=51 cells per condition).

**Figure 4-figure supplement 1.**
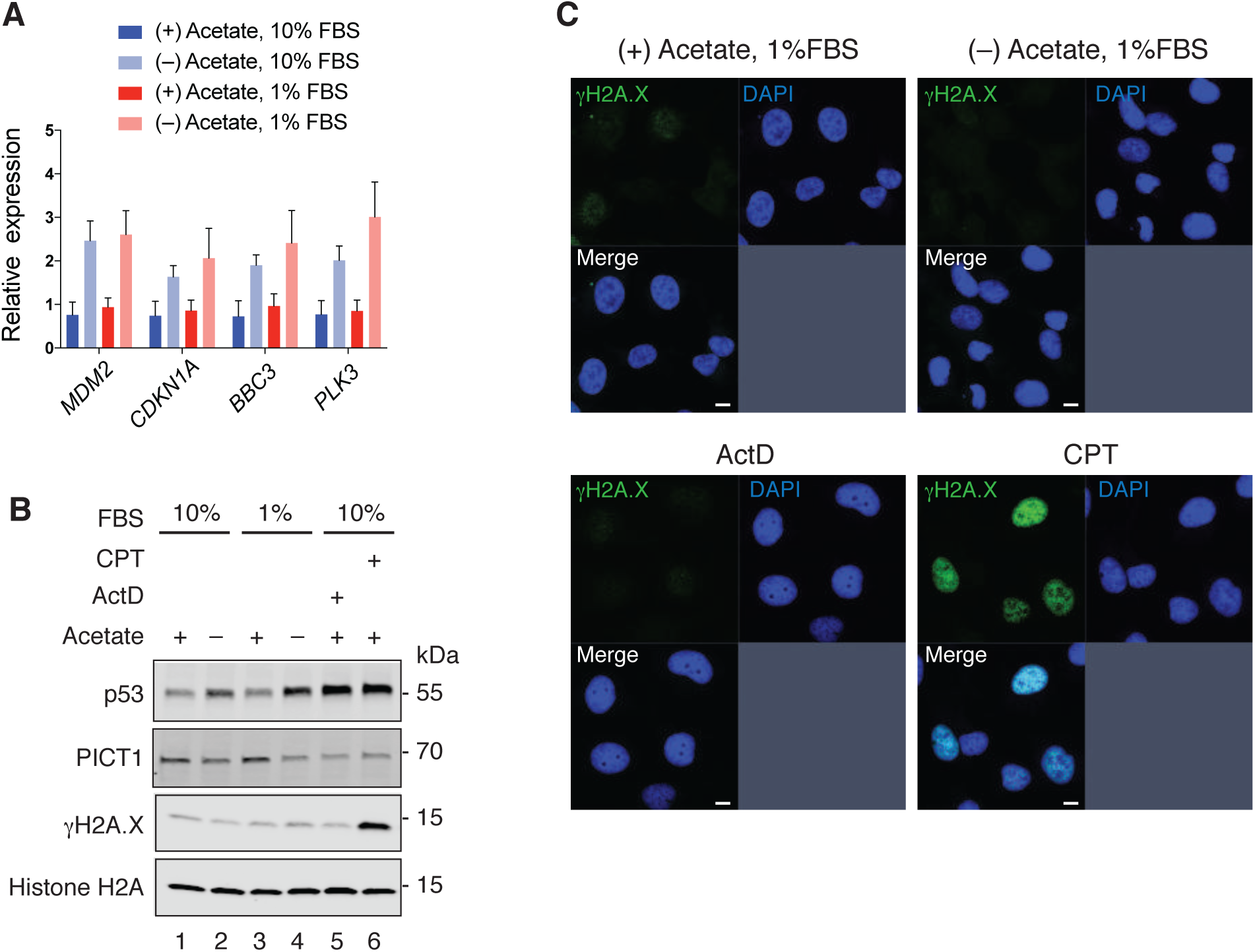
(A) Relative mRNA expressions for p53 target genes selected from the RNA sequencing in Table S2. Data are shown as mean ± S.D. (n=3 biological replicates). (B) Immunoblotting for p53, PICT1, *γ*H2A.X and Histone H2A in ASA-KO cells cultured in 10% or 1% FBS containing media with or without acetate for 4 hours, or treated with 5 nM ActD or 1 μM Camptothecin (CPT), a DNA damage inducer, for 4 hours. (C) Immunostaining for *γ*H2A.X in ASA-KO cells cultured in the same conditions as in (A).

**Figure 5-figure supplement 1.**
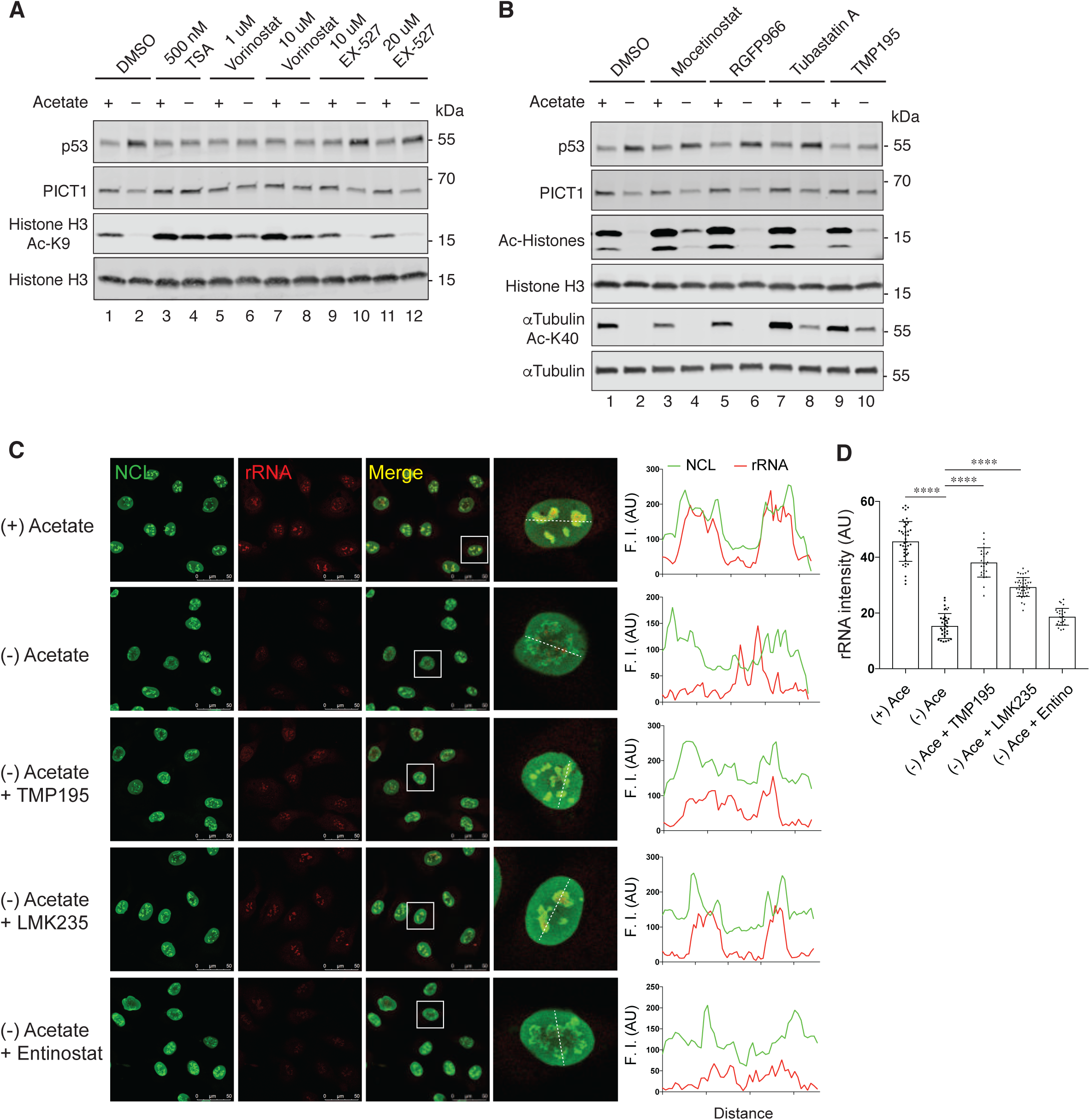
(A) Immunoblotting for the indicated proteins in ASA-KO cells cultured in 1% FBS containing media with or without acetate, and in the presence or absence of indicated HDAC inhibitors for 4 hours. (B) Immunoblotting for indicated proteins in ASA-KO cells cultured in 1% FBS containing media with or without acetate, and in the presence or absence of indicated HDAC inhibitors (10 μM) for 4 hours. (C) Immunostaining for NCL along with rRNA staining in ASA-KO cells cultured in 1% FBS containing meida with or without acetate and in the presence or absence of indicated HDAC inhibitors (50 μM TMP195, 10 μM LMK or 50 μM Entinostat) for 4 hours. Representative nuclear images (surrounded by a white square) are shown. Line profiles for indicated fluorescent intensities (F. I) determined along the white dashed lines are shown to the right. AU, arbitrary unit. (D) Quantification of the mean fluorescent intensity of the RNA signal per nucleus in (C). Data are shown as mean ± S.D. (n=29-53 cells per condition). ****P<0.0001 (One-way ANOVA followed by Tukey multiple comparisons test). AU, arbitrary unit.

**Figure 5-figure supplement 2.**
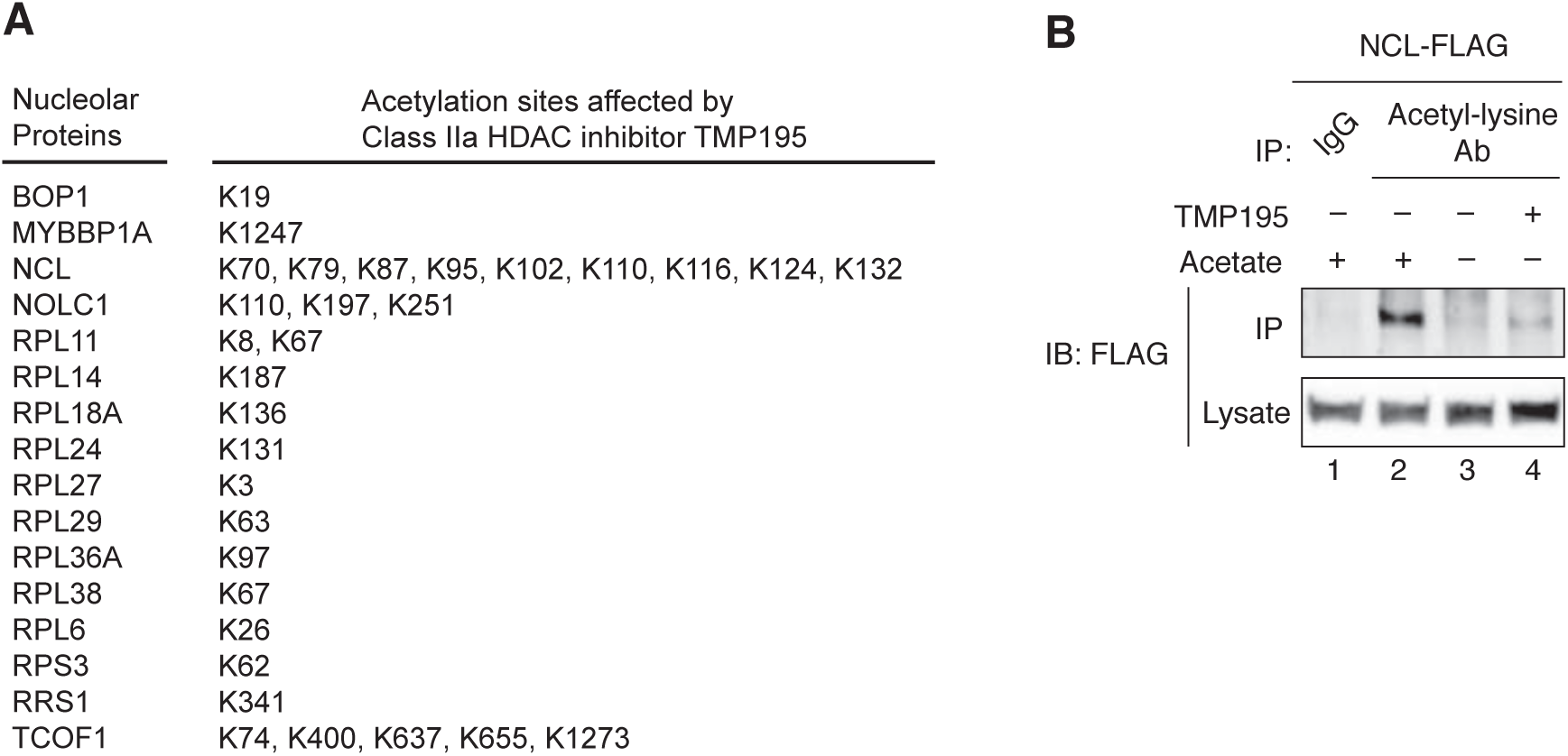
(A) Selected putative acetylated nucleolar proteins from Table S1 and their acetylation sites that are sensitive to TMP195 treatment. (B) Immunoprecipitation with anti-acetylated-lysine antibody and immunoblotting for NCL-3xFLAG in ASA-KO cells cultured in 1% FBS containing media with or without acetate and in the presence or absence of 50 μM TMP195 for 90 minutes.

**Figure 6-figure supplement 1.**
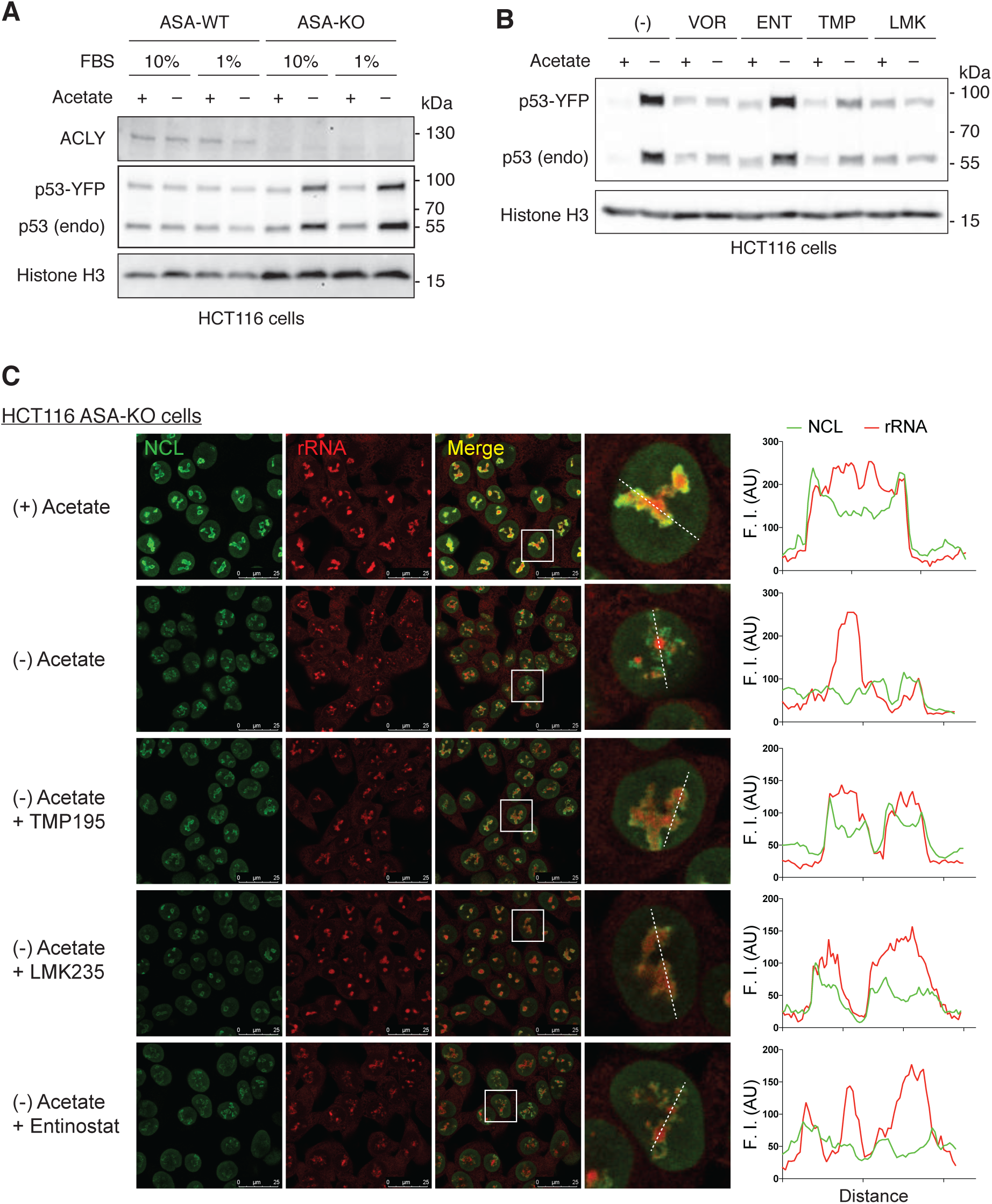
(A) Immunoblotting for p53 in HCT116 p53-YFP ASA-WT and ASA-KO cells cultured in 10% or 1% FBS containing media with or without acetate for 4 hours. Histone H3 is shown as a loading control. (B) Immunoblotting for the indicated proteins in HCT116 p53-YFP ASA-KO cells cultured in 1% FBS containing media with or without acetate, and in the presence or absence of indicated HDAC inhibitors for 4 hours. Following concentrations of HDAC inhibitors were used: 10 mM Vorinostat, 70 mM Entinostat, 50 mM TMP195 and 10 μM LMK235. (C) Immunostaining for NCL along with rRNA dye staining in ASA-KO cells cultured in 1% FBS containing media with or without acetate and in the presence or absence of indicated HDAC inhibitors (50 μM TMP195, 10 μM LMK or 50 μM Entinostat) for 4 hours. Magnified nuclear images (surrounded by a white square) are shown. Line profiles for indicated fluorescent intensities (F. I) determined along the white dashed lines are shown to the right.

